# Genome-wide absolute quantification of chromatin looping

**DOI:** 10.1101/2025.01.13.632736

**Authors:** James M. Jusuf, Simon Grosse-Holz, Michele Gabriele, Pia Mach, Ilya M. Flyamer, Christoph Zechner, Luca Giorgetti, Leonid A. Mirny, Anders S. Hansen

**Affiliations:** Department of Biological Engineering, Massachusetts Institute of Technology, Cambridge, MA 02139, USA; Gene Regulation Observatory, Broad Institute of MIT and Harvard, Cambridge, MA 02139, USA; The Novo Nordisk Foundation Center for Genomic Mechanisms of Disease, Broad Institute of MIT and Harvard, Cambridge, MA 02142, USA; Koch Institute for Integrative Cancer Research, Cambridge, MA 02139, USA; Center for Systems Biology Dresden, 01307 Dresden, Germany; Max-Planck Institute for the Physics of Complex Systems, 01187 Dresden, Germany; Friedrich Miescher Institute for Biomedical Research, 4065 Basel, Switzerland; University of Basel, 4001 Basel, Switzerland; Scuola Internazionale Superiori di Studi Avanzati, 34136 Trieste, Italy; Max Planck Institute of Molecular Cell Biology and Genetics, 01307 Dresden, Germany; Institute for Medical Engineering and Science, Massachusetts Institute of Technology, Cambridge, MA 02139, USA; Department of Physics, Massachusetts Institute of Technology, Cambridge, MA 02139, USA

**Author notes:** Division of Molecular Genetics and ONCODE Institute, Netherlands Cancer Institute, 1066CX Amsterdam, the Netherlands (Pia Mach).

## Abstract

3D genomics methods such as Hi-C and Micro-C have uncovered chromatin loops across the genome and linked these loops to gene regulation. However, these methods only measure 3D interaction probabilities on a relative scale. Here, we overcome this limitation by using live imaging data to calibrate Micro-C in mouse embryonic stem cells, thus obtaining absolute looping probabilities for 36,804 chromatin loops across the genome. We find that the looped state is generally rare, with a mean probability of 2.3% and a maximum of 26% across the quantified loops. On average, CTCF-CTCF loops are stronger than loops between cis-regulatory elements (3.2% vs. 1.1%). Our findings can be extended to human stem cells and differentiated cells under certain assumptions. Overall, we establish an approach for genome-wide absolute loop quantification and report that loops generally occur with low probabilities, generalizing recent live imaging results to the whole genome.

## INTRODUCTION

Chromatin looping, the process by which the genome folds to bring two regions into physical proximity, plays important roles in transcriptional regulation, DNA repair, genome integrity, DNA replication, and somatic recombination.^1–7^ Loops between enhancers and promoters are generally understood to facilitate transcription activation, while loops between CTCF sites, held together by the cohesin complex, mediate both insulation and facilitation by separating the genome into topologically associating domains (TADs).^2,8–10^ Disruption of both types of looping has been shown to cause aberrant gene regulation and lead to disease.^11–19^ Nevertheless, a general quantitative understanding of chromatin looping is currently lacking.^2,3^

Recent live imaging studies of a few individual chromatin loops have indicated that these loops are not permanent, stable structures but instead occur sporadically over time or across populations.^20–26^ Thus, the probability of loop formation represents an important biophysical quantity, which can be used to inform quantitative models of chromatin looping and its downstream effects.^27–34^ For example, it can be used to determine plausible parameter sets for kinetic models of enhancer-mediated transcriptional activation^30,34^ and DNA double-strand break repair,^35^ as well as polymer models of chromosome dynamics.^20,21,28,31– 33^ Rigorous quantification of looping probability is possible by applying Bayesian Inference of Looping Dynamics (BILD) on live imaging data with the necessary controls.^20^ However, since live imaging is experimentally intensive and low-throughput, only a few loops have been quantified, and the general trends of looping probability across the genome remain unknown.

Other methods have allowed chromatin looping to be observed in a genome-wide manner, but the inference of looping probabilities using these methods has been challenging. DNA fluorescence in situ hybridization (FISH) can be used to interrogate the 3D structure of chromatin in a high-throughput manner,^36–38^ but it is challenging to obtain precise looping probabilities from DNA FISH data due to localization uncertainty, the non-Markovian nature of chromatin loops, and the need for an arbitrary threshold.^2,39– 42^ Meanwhile, 3D genomics methods such as Hi-C and Micro-C^43–51^ also provide a genome-wide view of pairwise interactions, but their measurements are relative. This is an intrinsic limitation of all genomics methods, in which the signal is quantified from sequencing read counts, but calibration techniques such as spike-in normalization in RNA-seq^52^ and ChIP-seq^53–55^ have made absolute quantification of genomics experiments possible. Therefore, we sought to develop an absolute quantification method for 3D genomics experiments, allowing us to estimate absolute looping probabilities genome-wide.

Here, we combine Micro-C with BILD on live imaging data in mouse embryonic stem cells (mESCs) to achieve absolute quantifications for 36,804 loops across the genome. Our results indicate that loops occur with low probabilities, with a mean of 2.3%, median of 1.6%, and maximum of 26%. Looping probability varied between different classes of loops: CTCF/cohesin loops formed with higher probabilities than enhancer-promoter (E-P), promoter-promoter (P-P), and enhancer-enhancer (E-E) loops; more generally, absolute looping probability can be partially explained by the epigenomic features present at the loop anchors. Lastly, we demonstrated that under certain assumptions, our conclusion that the looped state is rare also extends to human cells. Overall, our findings indicate that rarity is a general property of all chromatin loops in mESCs and potentially mammalian cells in general.

## RESULTS

### Estimating absolute chromatin looping probabilities from Micro-C

We developed an approach to estimate the probabilities of looping, on an absolute scale, for all chromatin loops genome-wide from Micro-C data (**Fig. 1a**). Chromatin loops appear as focal enrichments (“dots”) in Micro-C maps.^45–48,56^ Since Micro-C maps quantify the degree of pairwise contacts, the strength of these dots is related to the underlying looping probability, i.e., the average fraction of alleles in which the loop exists at a given time. By measuring the dot strengths of loops whose absolute looping probabilities are known from live imaging and BILD^20^, we can obtain a calibration that allows us to estimate the absolute looping probability of any other loop in the genome from Micro-C alone (**Fig. 1a**).

**Figure 1.**
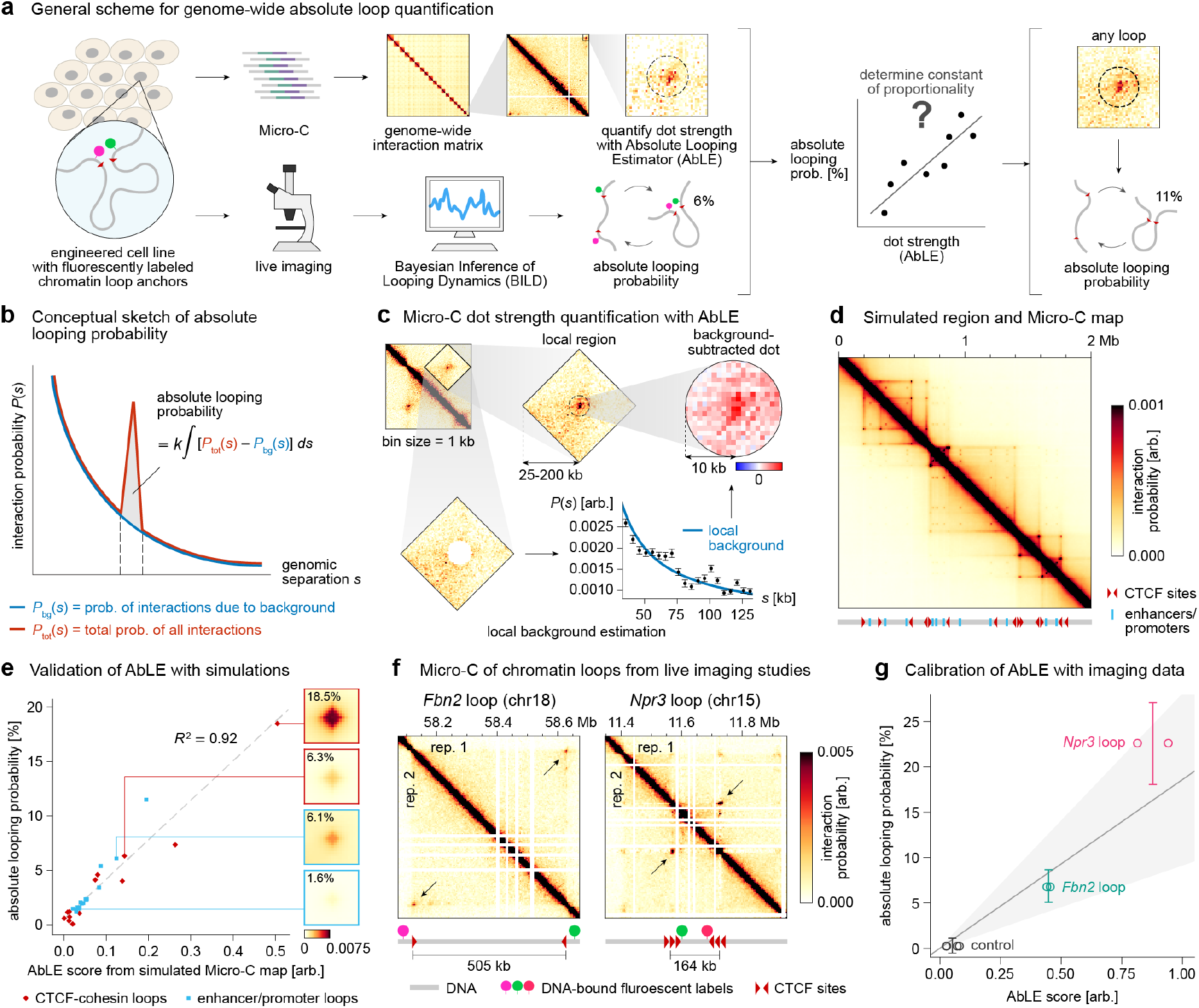
Overall scheme for genome-wide absolute quantification of chromatin loops from Micro-C. **a**, Flowchart describing genome-wide absolute loop quantification by calibrating Micro-C dot strength against loops measured in live imaging. **b**, Absolute looping probability is defined as the probability of interactions excluding the expected background. The background is primarily a function of genomic separation. **c**, AbLE quantifies Micro-C dot strength by summing the Micro-C signal originating from a chromatin loop excluding the local background, which is estimated in a separate step. The size of the local region, which can vary from 25-200 kb, is chosen to be proportional to 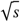 (see **Methods**). **d**, Diagram of simulated region with features (bottom) and simulated Micro-C map (top) from 3D polymer simulation. Micro-C bin size = 1 monomer, corresponding to 1 kb. Color scale is linear. **e**, Plot of ground-truth absolute looping probability from 3D polymer simulations vs. AbLE scores from the simulated Micro-C map. Gray dotted line indicates best-fit line of proportionality. Examples of simulated Micro-C dots corresponding to loops with various looping probabilities are shown on right. Micro-C color scale is linear. **f**, Micro-C dots corresponding to loops previously measured with live imaging; diagrams of engineered loci for live imaging shown below Micro-C maps. Micro-C bin size = 1 kb. Color scale is linear. Coordinates are given in terms of the modified genomes for the engineered cell lines (see **Fig. S9a**). **g**, Looping probability vs. AbLE score for *Fbn2* loop, *Npr3* loop, and control condition (*Fbn2* with depletion of RAD21). Gray line indicates best-fit line of proportionality; light gray shaded area indicates 95% confidence interval. Circles indicate AbLE scores of individual Micro-C replicates; error bars indicate bootstrapped 95% confidence intervals in BILD estimates.

To quantify looping, a precise definition of looping probability is required. An important consideration is that all pairs of DNA loci are subject to a background level of nonspecific interactions due to random polymer motion and actively extruding cohesins; the probability of these interactions is described by a *P*(*s*) curve, which is a function of the genomic separation *s* between the two loci (**Fig. 1b**). Loops are associated with a site-specific enrichment of interactions, presumably caused by mechanisms such as paused cohesin at CTCF sites and affinity-mediated interactions between transcription regulatory factors (**Fig. S1**). In this study, we explicitly define looping probability to only include the interactions above the background; this definition directly corresponds to what BILD quantifies in live-imaging data^20^. Therefore, the absolute looping probability is proportional to the sum of background-subtracted normalized Micro-C signal contributing to this enrichment, with an unknown scaling factor *k* (**Fig. 1b**, *k* ∫ [*P*_tot_(*s*)– *P*_bg_(*s*)] *ds*. Additionally, we note that Micro-C measures the fraction of alleles containing a loop at a single time point, whereas live imaging measures the fraction of time a loop is formed at a single allele (previously called *looped fraction*^20^). Here, we proceed by assuming that chromatin looping is an ergodic process, i.e., both quantities are equal, and we refer to them as the *looping probability*.

We developed a new method for quantifying Micro-C dot strength called AbLE (**Ab**solute **L**ooping **E**stimator) that sums the normalized Micro-C signal above background (**Fig. 1c**). Since the background *P*(*s*) curve is region-dependent, AbLE first estimates a local *P*(*s*) curve by fitting the values of the Micro-C map surrounding, but not including, the dot. The AbLE score is then calculated as the sum of background-subtracted values in a 10-kb circular region. AbLE is therefore designed to quantify the amount of Micro-C signal originating from looping events as previously defined.

To validate that AbLE measures looping probability, we performed 3D molecular dynamics simulations of a DNA polymer with CTCF/cohesin-mediated looping and affinity-mediated looping, the latter being a plausible mechanism for loops between enhancers and/or promoters^56,57^ (**Fig. 1d**). We observe a strong linear relationship (*R*^2^ = 0.92) between the AbLE scores from the simulated Micro-C map and the ground-truth absolute looping probabilities calculated directly from the individual molecular trajectories in the simulation (**Fig. 1e**). The linear relationship is robust to changes in the capture radius used to generate the simulated Micro-C maps and the interaction radius used to call ground-truth enhancer/promoter looping events^2^ (**Fig. S2**).

Next, we applied AbLE on experimental data. We performed deep Micro-C on two mESC cell lines (two replicates per cell line, ∼1.8 billion unique interactions per replicate, ∼7.4 billion unique interactions total; **Fig. S3a**), each containing a BILD-quantified loop: an endogenous 505-kb loop containing the *Fbn2* gene on chromosome 18^20^ and a synthetic 164-kb loop near the *Npr3* gene on chromosome 15 engineered to be extremely strong using triplets of high-occupancy CTCF sites^21^ (**Fig. 1f**). The Micro-C maps exhibited high reproducibility across replicates and cell lines (**Fig. S3b**). The empirically determined *P*(*s*) curves were approximately inversely proportional to the genomic separation *s* on average (**Fig. S3c**) but exhibited significant local variation (**Fig. S3d-f**), highlighting the need for the local background estimation step in AbLE. From our Micro-C data, we calculated the AbLE scores of the *Fbn2* and *Npr3* loops (**Fig. S4**) and compared them against their known looping probabilities from live imaging (**Fig. 1g**). As a negative control, we included the *Fbn2* loop after depletion of cohesin subunit RAD21^20^, which exhibited near-zero looping probability in both Micro-C and live imaging data as expected. Fitting a line through the origin and the *Fbn2* and *Npr3* points resulted in a best-fit slope of *k* = 0.186 ± 0.047. In summary, we developed AbLE to measure the strength of loops from Micro-C and calibrated it against loops with known absolute looping probabilities from live imaging.

### Global analysis of chromatin looping probabilities

Having developed an approach to perform absolute 3D genomics, we sought to apply it to understand general trends in looping probability across the genome. To call loops across the genome with high confidence, we first generated an extremely deep mESC Micro-C map (15.6 billion unique interactions) by merging our Micro-C data with those from past studies in mESCs^45,46,48,58^ (**Fig. 2a**, datasets listed in **Table S1**), which we verified to be highly similar (**Fig. S5a**). The superior depth of this Micro-C map resulted in a greatly improved signal-to-noise ratio over maps from individual samples (**Fig. S5b**). From the merged Micro-C map, we identified 153,658 loops across the genome using the *Mustache* loop caller^59^ (**Table S2**). After removing loops with insufficient reads, missing data nearby, or interference from nearby loops or the diagonal of the Micro-C map (**Fig. 2a**, see **Methods**), we obtained a final set of 36,804 loops for quantification.

**Figure 2.**
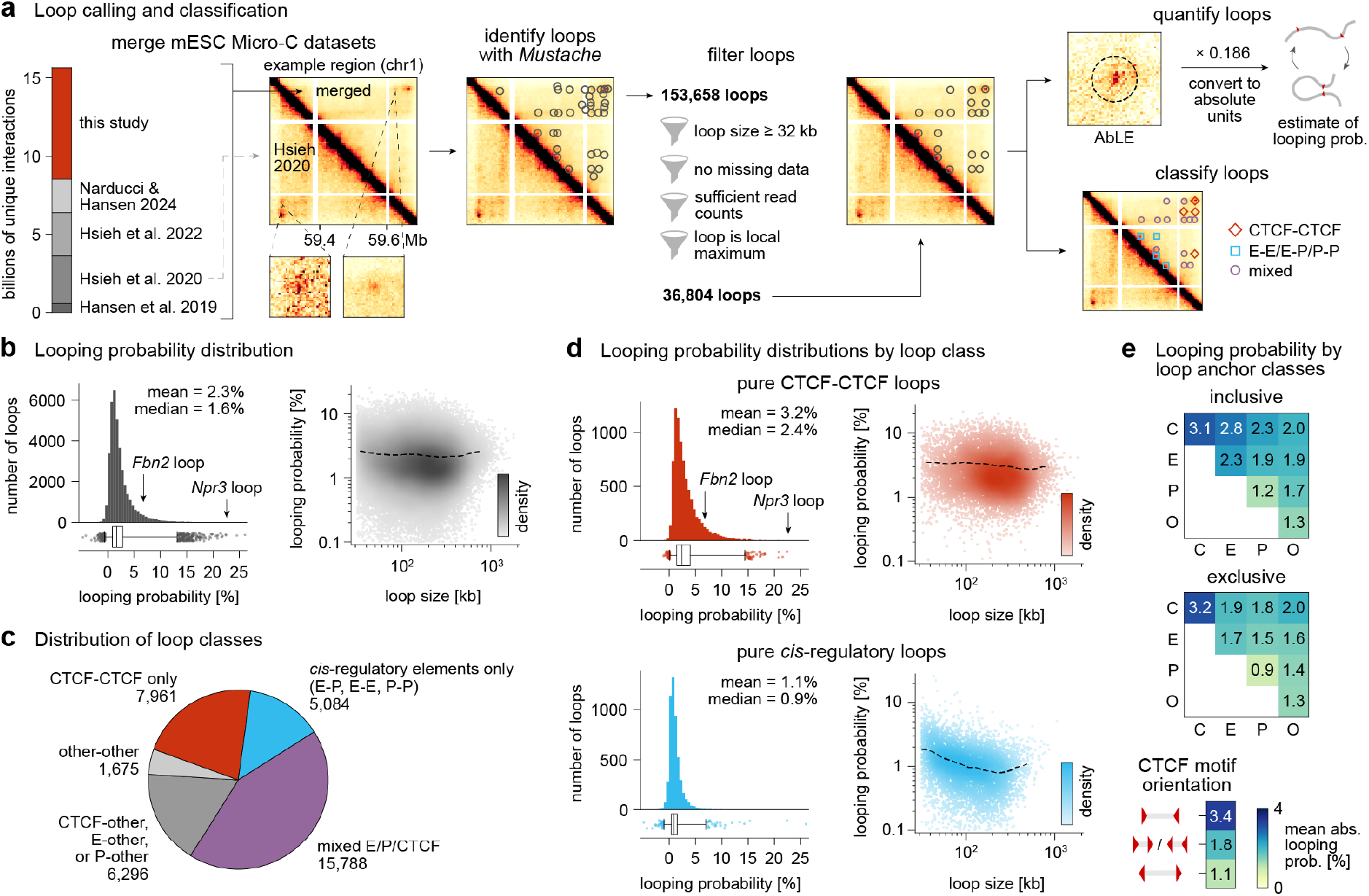
Global analysis of chromatin looping probabilities. **a**, Flowchart of methods used to perform global analysis of chromatin looping probabilities. Five mESC Micro-C datasets were combined to form a merged dataset with superior signal-to-noise ratio compared to the deepest dataset previously available.^45^ Then, loops were called using *Mustache* and filtered for quantifiability. Filtered loops were quantified and classified for further analysis. **b**, Distribution of looping probabilities and relationship between probability vs. size among 36,804 filtered loops. In the box plot, whiskers extend to 0.5^th^ and 99.5^th^ percentiles; all outliers beyond the whiskers are plotted as individual points. **c**, Distribution of loop classes as determined by presence of CTCF/cohesin, enhancers, or promoters at loop anchors. “Other” refers to anchors that did not overlap any of these features. **d**, Distributions of looping probabilities and relationship between probability vs. size among pure CTCF-CTCF loops and pure *cis*-regulatory loops. **e**, Mean looping probability for all possible combinations of loop anchor classes. Under the “inclusive” definition, a loop anchor is said to be a CTCF- and cohesin-bound site, enhancer, or promoter if that feature is present, regardless of whether other features are present. Under the “exclusive” definition, a loop anchor is said to have a feature if only that feature is present; loops with two or more features at either anchor are omitted from the analysis. C = CTCF/cohesin, E = enhancer, P = promoter, and O = other. Among all CTCF-CTCF loops (defined with the inclusive definition), the effect of CTCF motif orientation is also analyzed.

To estimate their absolute looping probabilities, we quantified these loops with AbLE in each of our four replicates of Micro-C independently, averaged the scores across replicates, and multiplied the scores by the previously obtained calibration factor, *k* = 0.186 (**Fig. 1g**; quantifications in **Table S3**). The AbLE scores showed high agreement across replicates and cell lines (**Fig. S5c**). As a negative control, we verified that on average, the AbLE score measured at random locations not corresponding to loops was equal to zero within statistical error (**Fig. S5d**). Further cross-validation with live imaging data was not possible due to the absence of other live imaging experiments containing the necessary controls for BILD, so we instead turned to DNA FISH studies.^60,61^ We quantified E-P looping probabilities of key E-P pairs in mESCs: 19.8% for *Sox2* (integrating the entire ∼7-kb *Sox2* control region (SCR)^62^), 12.7% for *Mycn*, 5.8% for *Klf4*, and 5.3% for *Nanog* (**Fig. S5e**). Despite significant differences between our method and DNA FISH and the exact definitions of the genomic regions involved, the results generally agree within a factor of two (**Fig. S5f**). More generally, in our experimental data, we estimate the fractional uncertainty in our looping estimates to be ∼30-40%, which arises from uncertainty in the AbLE score (affected by biological noise and Micro-C read count noise) and uncertainty in the scaling factor *k* (**Fig. S5g**). For the weakest loops (<0.5%), the fractional uncertainty increases to beyond 40%. Uncertainty among very weak loops results in a small fraction of loops with an apparent negative looping probability; however, we emphasize that this is a technical artifact due to noise.

Having cross-validated our approach, we turned to global trends among the 36,804 quantified loops (**Fig. 2b**). Overall, looping probabilities ranged from much less than 1% to 26%, with a mean of 2.3% and a median of 1.6% (**Fig. 2b**). Thus, the well-known SCR-*Sox2* E-P loop is among the strongest in the genome (**Fig. S5b**,**e**). We also note a loop selection bias: had we been able to quantify all 153,658 loops, which includes many very weak loops, the mean and median would likely have been lower. Conversely, had we quantified only the strongest loops, the mean and median looping probability would have been higher. Nevertheless, our results clearly show that in mESCs, all loops only exist in a fraction of cells at a given time, and most loops occur very rarely.

We also compared our absolute loop quantifications against a commonly-used metric of relative loop strength, the logarithm of observed signal divided by expected signal. The two quantities were generally positively correlated, but the log-ratio metric was disproportionately greater for larger loops, likely due to the very low expected background (**Fig. S5h**).

Next, we classified loops based on whether their anchors were near CTCF- and cohesin-bound sites, enhancers, or promoters. CTCF- and cohesin-bound sites were identified using CTCF and SMC1A ChIP-seq peaks and CTCF binding motifs, promoters were defined as regions ±2 kb from transcription start sites,^63^ and enhancers were defined as regions of overlapping H3K4me1 and H3K27ac ChIP-seq peaks. We detected 7,961 pure CTCF-CTCF loops (containing only CTCF/cohesin at the anchors), 5,084 pure *cis*-regulatory loops (containing only enhancers and/or promoters at the anchors), 15,788 mixed loops (containing both CTCF/cohesin and *cis*-regulatory elements at the anchors), and 7,971 other combinations (**Fig. 2c;** classifications in **Table S3**). All loop classes exhibited a wide range of looping probabilities (diverse examples shown in **Fig. S6a**), though CTCF-CTCF loops were stronger than *cis*-regulatory loops on average, with a mean looping probability of 3.2% versus 1.1% (**Fig. 2d**). Also, CTCF-CTCF looping probabilities showed no dependence on loop size, whereas *cis*-regulatory looping probabilities showed a weak negative correlation with loop size (**Fig. 2d**).

Because many loops were anchored by a mixture of CTCF sites and *cis*-regulatory elements, we next calculated the mean looping probabilities for all possible combinations of loop anchor classes (**Fig. 2e**). The highest mean looping probability was found among pure CTCF-CTCF loops (3.2%), while the lowest was found among pure P-P loops (0.9%). The finding that P-P loops were weaker than E-P and E-E loops differs from prior work^45,46^ and is primarily driven by many weak P-P loops newly identified in this study (**Fig. S6b**). For loops anchored by CTCF sites, the orientation of the CTCF binding motifs strongly affected absolute looping probability, with convergent motifs forming the strongest loops as expected^43,64–67^ (**Fig. 2e**).

Having observed the differences in chromatin looping probabilities between known loop classes, we next sought to characterize the quantitative relationship between looping probabilities and a broader set of epigenomic features at the loop anchors in an unsupervised manner.

### Chromatin looping probabilities are correlated with epigenomic features at loop anchors

We examined the degree to which absolute looping probability is governed by epigenomic features at the loop anchors such as histone modifications, looping factors, and transcription factor (TF) binding (determined by ChIP-seq), chromatin accessibility (ATAC-seq), and active transcription (GRO-seq). In total, we considered 43 features from publicly available datasets (listed in **Table S4**), quantifying the signal from each feature within 5-kb windows centered on the left and right anchors of each loop (**Fig. 3a**; quantifications in **Table S5**). The feature strengths at both loop anchors affected the mean probability of looping; for example, the strengths of CTCF and the cohesin subunits SMC1A and RAD21 at loop anchors were positively correlated with looping probability (**Fig. 3b**). The variation of these features spanned a wide range of absolute looping probabilities; for example, loops with RAD21 binding in the highest decile at both loop anchors had a mean looping probability of 4.9% (90th percentile among all loops), whereas loops with binding in the lowest decile at both anchors had a mean looping probability of 1.5% (46th percentile among all loops). Other features such as H3K27ac, H3K4me3, and RNA Pol II were negatively correlated with looping probability (**Fig. 3b**). Similar heatmaps for the remaining 37 features are shown in **Fig. S7**. In general, our results showed that the probability of a loop is correlated with the strengths of epigenomic features at its anchors.

**Figure 3.**
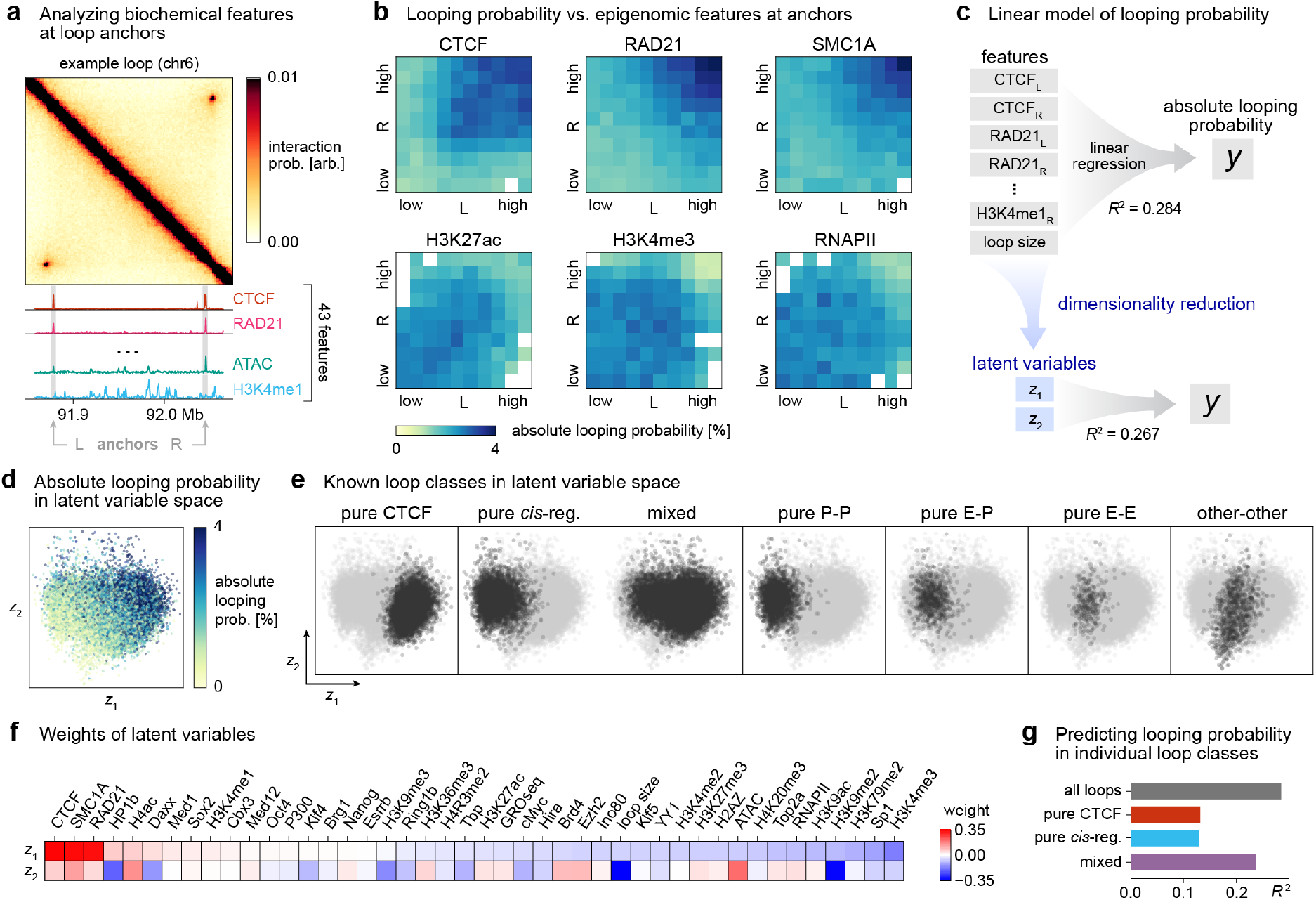
Chromatin looping probabilities are correlated with epigenomic features at loop anchors. **a**, Epigenomic features at each anchor of each loop are quantified by summing the values from the associated assay (ChIP-seq, ATAC-seq, or GRO-seq) in a 5-kb window centered on the anchor. Micro-C and select tracks are shown for an example loop. Micro-C bin size = 1 kb. Color scale is linear. **b**, Looping probability vs. feature strengths at anchors for six ChIP-seq features. *x*- and *y*-values represent ChIP signals at left (L) and right (R) anchors and are binned into deciles; color scale represents the mean absolute looping probability within each bin. Bins in which the standard error of the mean (*σ*/n) exceeds 0.2 are colored white due to their high uncertainty. **c**, Schematic of linear models for predicting looping probability from epigenomic features. Top: OLS linear regression model with 87 regressors. Bottom: dimensionality-reduced model with 2 latent variables. Models are trained on a subset containing 80% of loops; *R*^2^ values are calculated the remaining 20% of data. **d**, Scatter plot of all filtered loops in latent variable space, colored by absolute looping probability. **e**, Scatter plots of loops in latent variable space, with loops in specific classes highlighted in black and other loops colored gray. **f**, Weight vectors defining the transformation to latent variable space. Due to symmetry between the left and right anchors, the weights for each feature represents both the weight for the left and right anchors (e.g., “CTCF” represents the weights for regressors CTCF_L_ and CTCF_R_). **g**, Goodness-of-fit *R*^2^ of OLS models for predicting absolute looping probability from all 87 regressors, within loop classes individually. Models are trained on a subset containing 80% of loops; *R*^2^ values are calculated the remaining 20% of data.

We next asked how well absolute looping probability could be predicted from the strengths of epigenomic features at the anchors. We employed a simple linear model with all the strengths of all 43 features at left and right anchors as inputs (log-transformed and standardized; see **Methods** for details). Furthermore, since loop size was observed to vary with looping probability for *cis*-regulatory loops (**Fig. 2d**), we included loop size as an additional regressor, resulting in 87 regressors in total. We fitted the model using ordinary least squares (OLS) on a randomly selected training dataset consisting of 23,858 filtered loops; the model predicted absolute looping probability in a testing dataset of 5,964 loops with *R*^2^ = 0.284 (**Fig. 3c**). Thus, epigenomic features at loop anchors alone are somewhat predictive of absolute looping probability, but there is still significant unexplained variability. This suggests that looping probability is highly dependent on other contextual factors beyond what exists at the loop anchors; for example, these could be nearby CTCF sites, TADs, or synergy/competition between loci that participate in multiple loops.

Since epigenomic features are often correlated, we asked what specific combinations of features were most predictive of absolute looping probability. We performed dimensionality reduction on the features using projection to latent structures (PLS). By taking the first four PLS components and applying a varimax rotation to simplify the interpretation of the results^68^, we found two linear combinations of the original regressors that were sufficient to predict absolute looping probability with *R*^2^ = 0.267, performing nearly as well as the original OLS model (**Fig. 3c**). We designate these linear combinations as latent variables *z*_1_ and *z*_2_.

Plotting the loops in the latent variable space, we observe that higher values of *z*_1_ and *z*_2_ are associated with higher looping probabilities (**Fig. 3d**). Furthermore, the known loop classes formed distinct clusters in the latent variable space, primarily separating along the *z*_1_-axis (**Fig. 3e**). We also considered the weights of the original features in the latent variables: *z*_1_ contains strong positive weights for CTCF, SMC1A, and RAD21 and weak negative weights for many histone modifications, ATAC-seq, TFs, and other features associated with transcription regulation (**Fig. 3f**). This result explains the separation of CTCF/cohesin-bound vs. *cis*-regulatory loops along *z*_1_ and is consistent with our result that CTCF/cohesin-bound loops form with higher probabilities (**Fig. 2d**). The next component *z*_2_ contained strong positive weights for H4ac and ATAC-seq and strong negative weights for H3K9me2, HP1b, and loop size. This suggests that some signatures of active enhancers and promoters are associated with higher looping probability, whereas heterochromatin and larger loop sizes are negatively associated with looping probability.

Lastly, we asked whether looping probabilities could be predicted within the individual loop classes: pure CTCF-CTCF loops, pure *cis*-regulatory loops, and mixed loops. For each class of loops, we trained a linear model using OLS employing all 87 features without dimensionality reduction. On mixed loops, the model achieved *R*^2^ = 0.236, whereas the *R*^2^ values for pure CTCF-CTCF loops and pure *cis*-regulatory loops were 0.131 and 0.129, respectively (**Fig. 3g**). We conclude that the predictability of looping probability from epigenomic features at loop anchors primarily arises from the classification of loops into the two primary mechanisms (CTCF loops and E-P/P-P/E-E loops), and predictability decreases significantly when only considering loops arising from a single mechanism.

### Human dataset exhibits similar trends in chromatin looping

Next, we sought to test the generality of our mESC findings in human cells. Although coupled Micro-C with the necessary controls and live imaging data in human cells is not available, high-resolution Micro-C data is available in H1 hESCs and human foreskin fibroblasts (HFFs).^47^ These data were generated using the same experimental protocol as the mESC data in this study. We again called loops in these Micro-C datasets using *Mustache*^*59*^ and filtered them for quantifiability to obtain 19,763 loops in hESCs and 40,550 loops in HFFs (**Fig. 4a**, filtered loops in **Table S6**). To obtain rough estimates of the absolute looping probabilities, we calculated the AbLE scores of these loops and multiplied them by *k* = 0.186, applying the conversion factor from our mESC calibration (**Fig. 1e**). Since the same Micro-C experimental procedure was used to generate the mouse and human data, the values in the Micro-C matrices likely exist on a similar scale. Indeed, the *P*(*s*) curves of the mESC, hESC, and HFF data are very similar (**Fig. S8**). Therefore, we proceeded to apply our mESC conversion factor to the human data, though we stress that the uncertainty in our absolute looping probability estimates is substantially higher and that the results should be only considered as rough ballpark figures. Under these assumptions, we found that the looping probabilities in human cells had a similar distribution to those of mESCs, also occurring with low probabilities, with the majority being less than 10% (**Fig. 4b**, loop quantifications in **Table S6**). The mean looping probability was 4.2% in hESCs and 3.8% in HFFs. Among the loops detected, 11,391 loops were shared among both cell types. Among these loops, the mean looping probability was 5.0% in hESCs vs. 6.5% in HFFs (**Fig. 4b**), and 67% of loops had a higher looping probability in HFFs than in hESCs (**Fig. 4c**). Using publicly available ChIP-seq data to classify CTCF-CTCF loops, we found that CTCF-CTCF loops had higher than average looping probabilities, generalizing the result from mESCs (**Fig. 4b**). These results suggest that the low probabilities with which chromatin loops occur may be a general property of mouse and human stem cells and differentiated cells, and some differentiated cells may have slightly stronger loops than embryonic stem cells.

**Figure 4.**
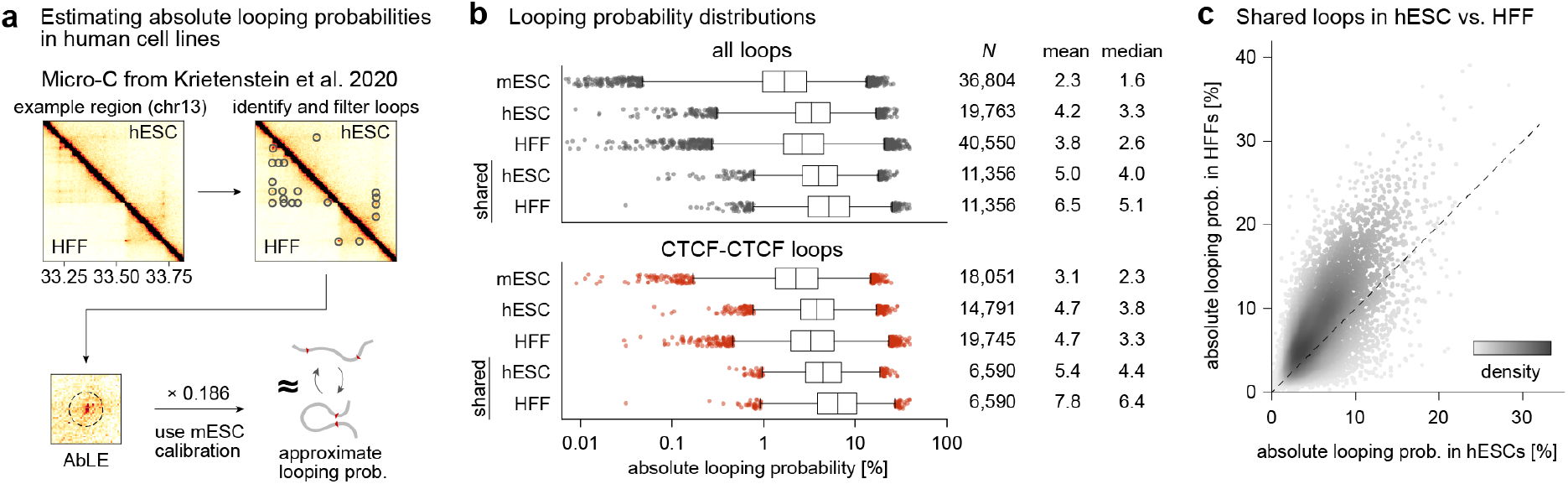
Human embryonic stem cells (hESCs) and human foreskin fibroblast cells (HFFs) exhibit similar chromatin looping trends as mESCs. **a**, Flowchart of method to estimate absolute looping probabilities in hESC and HFF cells. In each cell type, loops are called from Micro-C maps using *Mustache*, filtered for quantifiability, and quantified using AbLE. Quantifications are converted to absolute units using the mESC calibration. **b**, Looping probability distributions and statistics for mESC loops (**Fig. 2b,d**) hESC loops, HFF loops, and shared loops (loops common to both hESCs and HFFs) in hESCs and HFFs for all loops and for CTCF-CTCF loops. Box plot whiskers extend to 0.5^th^ and 99.5^th^ percentiles; outliers beyond the whiskers are plotted as individual points. Due to their high uncertainty, some loops with extremely low probabilities (below 10^−3^-10^−4^) are not shown, but these account for less than 3% of loops in each category. **c**, Scatter plot of looping probabilities of shared loops in HFFs vs. hESCs. Dashed line indicates the line *x* = *y*.

## DISCUSSION

In this study, we showed that absolute quantification of chromatin loops can be achieved by integrating deep Micro-C data quantified with AbLE coupled with live imaging data quantified with BILD^20^. Using this approach, we estimated absolute looping probabilities for 36,804 loops in mESCs and found that loops generally form with low probabilities, with a mean of 2.3% and maximum of 26% (**Fig. 2b**).

A major goal of the 4D nucleome project has been to integrate genomics and imaging data.^69^ Our results demonstrate that quantitatively integrating sequencing-based 3D interaction data and imaging data is feasible, provided the quantities being measured are consistent and rigorously defined. We acknowledge that the constant of proportionality that converts AbLE scores from Micro-C into absolute units cannot be derived theoretically. However, we show that AbLE is linearly correlated with absolute looping regardless of the interaction radius used in the 3D polymer simulations (**Fig. S2**). Therefore, determining the constant of proportionality empirically is sufficient to bridge genomics and imaging, even though the underlying physical parameters remain unknown.

What are the implications of low looping probabilities? Since Micro-C is a population-averaged snapshot at the time of the experiment, two interpretations are possible: either that loops occur infrequently in all cells, or that loops are permanent and stable in a certain subpopulation of cells and nonexistent in others. The significant variability needed for the latter explanation is unlikely in homogeneous cell populations. Therefore, our results support the notion that loops are short-lived, constantly forming and disappearing in single cells, generalizing the findings of recent live imaging studies of two specific loops.^20,21^

Our results showed that *cis*-regulatory loops, including E-P loops, occur with lower probabilities than CTCF-CTCF loops on average (**Fig. 2d,e**). This result agrees with previous studies.^1,6,45-47,56,57^ Considering the absolute numbers revealed in this study, the probabilities for *cis*-regulatory loops are strikingly low, only ∼1-2% on average (**Fig. 2d,e**). While several imaging studies have focused on the SCR-*Sox2* E-P loop due to its high biological interest,^23,26,60,61^ this loop is one of the strongest in the genome. Beyond *Sox2* and other classical examples (**Fig. S5e**,**f**), the vast majority of *cis*-regulatory loops across the genome are generally extremely weak (**Fig 2d**,**e**). Given a typical mESC cell cycle of ∼600 minutes and looping probabilities of ∼1-2%, most *cis*-regulatory elements loop for a total ∼6-12 min per cell cycle beyond background chromosomal interactions. Thus, unless *cis*-regulatory loops form extremely infrequently, our results suggest that the lifetime of *cis*-regulatory loops is likely shorter than the ∼20 min lifetime of CTCF loops.^20,21^ The predominance of these low-probability loops suggests that E-P interactions may not require sustained contact to perform their transcription regulatory function, but instead form more frequently and very transiently.

Finally, we state the limitations and future directions of our work. First, we are currently limited to only two data points in our calibration (**Fig. 1g**), and our absolute loop quantifications are therefore associated with high uncertainty and should only be considered estimates. We look forward to extending our approach by incorporating more live imaging data as they become available. Another second limitation is that we are only able to report on the background-subtracted probability of looping (**Fig. 1b**) because this is what can be rigorously quantified using BILD.^20^ For some functional interactions such as E-P looping, transcription likely depends on total looping probability (without background subtraction). Third, to rigorously apply our absolute 3D genomics approach to other species and cell types, parallel live imaging and Micro-C experiments are required. Nevertheless, subject to the strong assumption of a constant calibration factor for Micro-C data which is aided by the robustness to capture radius (**Fig. S2**), our approach can approximately be applied to any cell type for which Micro-C data exists (**Fig. 4**). Fourth, it is impossible to determine loop lifetimes from fixed-cell data such as Micro-C and DNA FISH. Live imaging data will therefore always be required to understand the underlying dynamics that contribute to the low chromatin looping probabilities we observe.

In summary, by integrating live imaging with Micro-C, we present an absolute 3D genomics approach that reveals genome-wide absolute looping probabilities, finding that nearly all loops occur with low probabilities.

## Acknowledgements

We thank J. H. Yang, M. K. Huseyin, and A. A. Galitsyna for discussions on computational analyses. We also thank S. Kim, M. Nagano, D. N. Narducci, V. Ramanathan, J. Toppen, and J. H. Yang for insightful comments on the manuscript. A.S.H. acknowledges funding support from the NIH (DP2GM140938, R33CA257878, R01EB035127, UM1HG011536, R01CA300848, R03OD038390), an NSF CAREER award (2337728), the Gene Regulation Observatory of the Broad Institute of MIT and Harvard, the Novo Nordisk Foundation (NNF21SA0072102), a Pew-Stewart Scholar for Cancer Research award, the Mathers Foundation, and an RSC award from the MIT Westaway Fund. This work was supported by the Bridge Project, a partnership between the Koch Institute for Integrative Cancer Research at MIT and the Dana-Farber/Harvard Cancer Center. We thank the MIT Koch Institute’s Robert A. Swanson (1969) Biotechnology Center for technical support, specifically the Integrated Genomics and Bioinformatics Core and MIT BioMicroCenter, and this work was supported in part by the Koch Institute Support (core) Grant P30-CA14051 from the National Cancer Institute. L.M. acknowledges funding support from the NIH NIGMS (R01GM114190) and NSF (awards 2044895 and 2210558). L.M. is a Simons Investigator of Simons Foundation. We also thank the Walk-Up Sequencing services of the Broad Institute of MIT and Harvard. P.M. is supported by the Marie Skłodowska-Curie Innovative Training Network grant (no. 813327, “ChromDesign”).

## Data Availability

The data generated in this study can be found at NCBI Gene Expression Omnibus under accession number GSE286495. The code associated with this study can be found at https://github.com/ahansenlab/AbsLoopQuant_analysis_code.

