## Supplementary Information for "Genome-wide absolute quantification of chromatin looping"

#### Contents

- Supplementary Figures & Figure Legends
- Materials and Methods
- Supplementary Tables

### SUPPLEMENTARY FIGURES & FIGURE LEGENDS

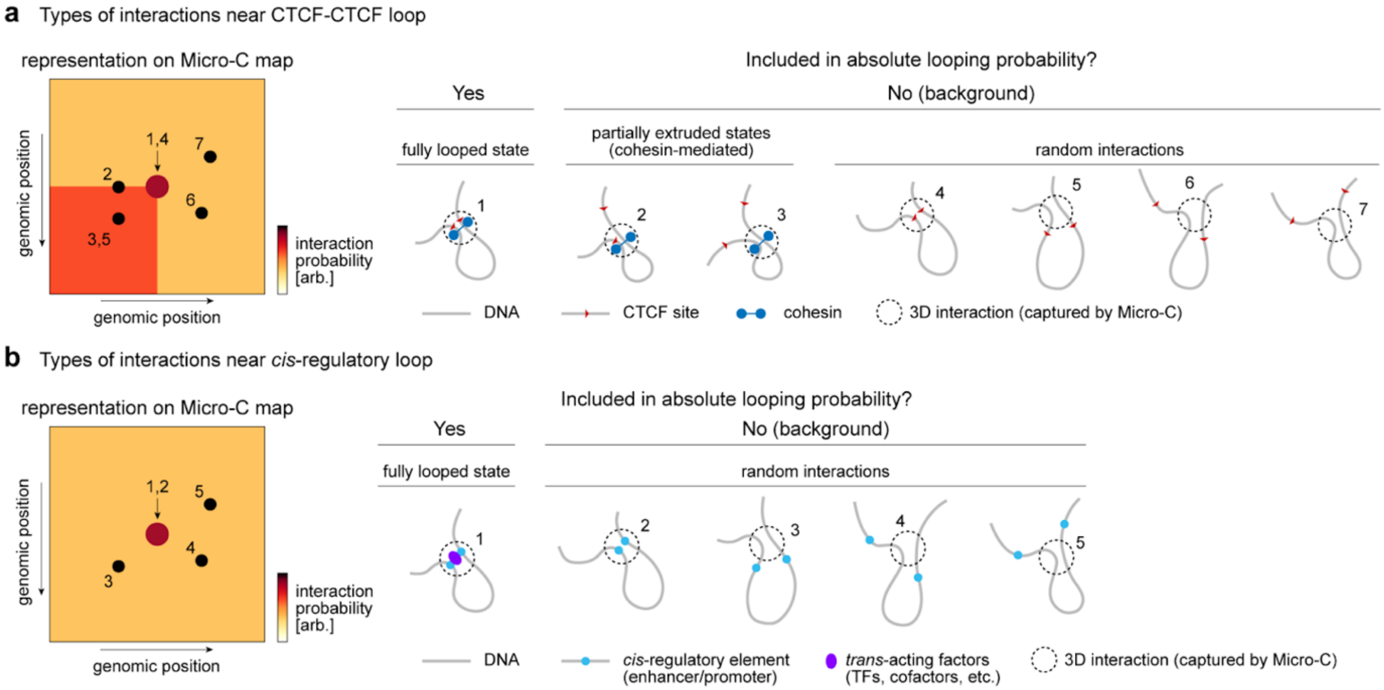


**Figure S1. Explanation of the scope of absolute looping probability for CTCF-CTCF and *cis*-regulatory loops. a,** For a CTCF-CTCF loop, we define a loop as an interaction between the two anchors mediated by a complex of CTCF and cohesin (1). Other interactions may result in reads on the Micro-C map nearby or at the loop, due to the activity of passing cohesin (2-3) or random polymer motion (4-7), but are excluded from the calculation of absolute looping probability. **b,** Similar to panel c, but for *cis*-regulatory loops.

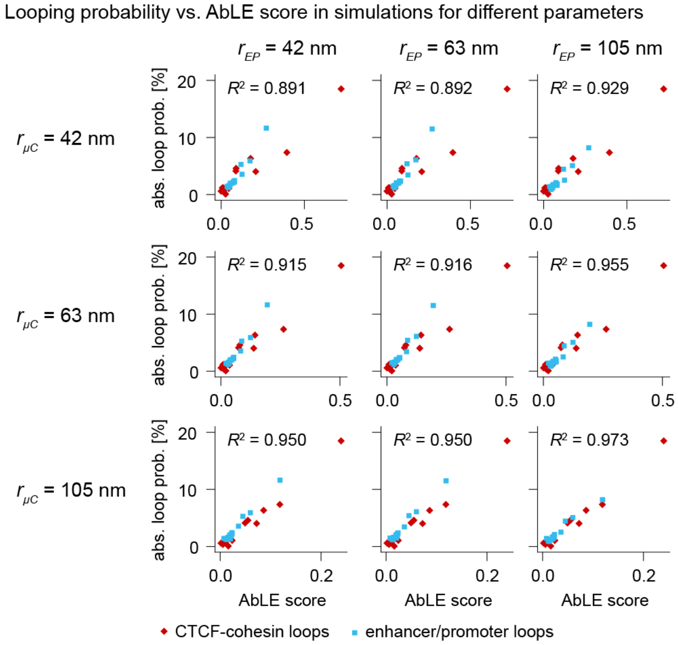


**Figure S2. Robustness of linear trend between AbLE and looping probability from simulated data.** Absolute looping probability (ground truth) vs. AbLE score in simulations as in **Fig. 1e**, shown for different values of the Micro-C capture radius ($r_{\mu C}$) and the radius used to declare ground-truth enhancer/promoter contacts ($r_{EP}$).

**a** Micro-C statistics

| | *Fbn2* cell line | | *Npr3* cell line | |
|---|---|---|---|---|
| | **replicate 1** | **replicate 2** | **replicate 1** | **replicate 2** |
| **Sequenced pairs** | 3,200,680,310 | 3,126,280,669 | 3,220,699,369 | 3,303,701,523 |
| **Mapped pairs** | 2,081,658,192 | 2,079,903,306 | 2,160,474,690 | 2,207,982,526 |
| **Unique pairs** | 1,832,408,206 | 1,811,887,747 | 1,873,343,139 | 1,910,932,534 |
| **Fraction of mapped pairs unique** | 88% | 87% | 87% | 87% |

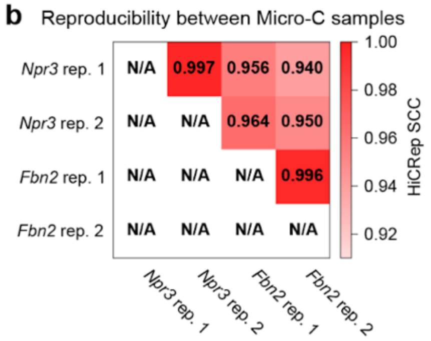


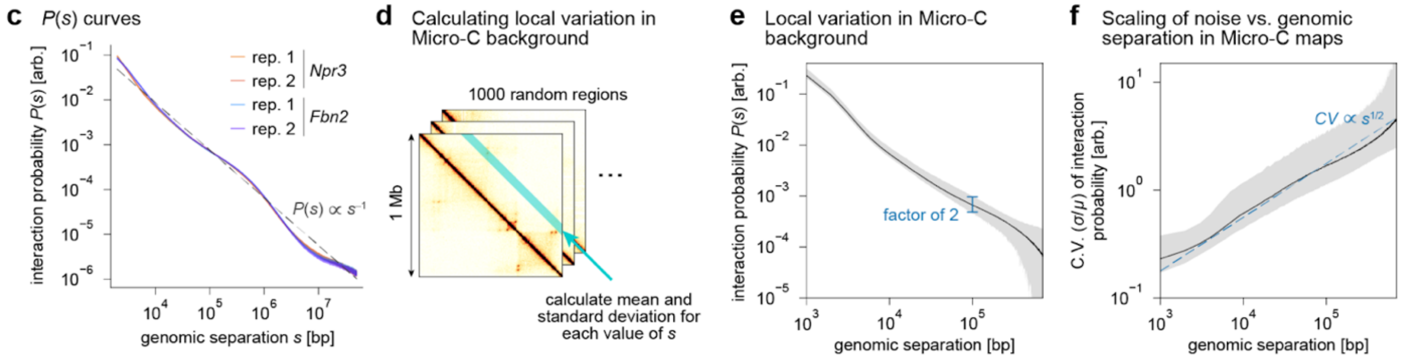


**Figure S3. Statistics and technical analyses of Micro-C data. a,** Statistics of Micro-C experiments performed for this study on the *Fbn2* cell line[20] and *Npr3* cell line[21]. **b,** Reproducibility between Micro-C replicates as quantified by HiCRep[70] stratum-adjusted correlation coefficient (SCC) at 100-kb resolution. **c,** Genome-wide averaged $P(s)$ curves calculated from Micro-C maps at 1-kb resolution. Gray dotted line shows the approximate power-law relationship between $P(s)$ and $s$. **d,** Method used to calculate statistics for panels e and f; the mean and standard deviations of all the matrix elements at separation $s$ are calculated for 1000 random 1 Mb regions. **e,** Aggregate of local $P(s)$ curves calculated on 1000 random regions genome-wide. The black line represents the mean; the shaded gray area represents the range between the 2.5$^{th}$ and 97.5$^{th}$ percentiles of $P(s)$ for each value of $s$. For a typical loop size of $10^5$ bp, there is a two-fold difference between the 2.5$^{th}$- and 97.5$^{th}$-percentile values of $P(s)$. **f,** Coefficient of variation (C.V.) of values in Micro-C map vs. genomic separation, within each 1 Mb region. The black line represents the mean; the shaded gray area represents the range between the 2.5$^{th}$ and 97.5$^{th}$ percentiles of C.V., for each value of $s$. Blue dotted line shows the approximate power-law relationship between C.V. and $s$.

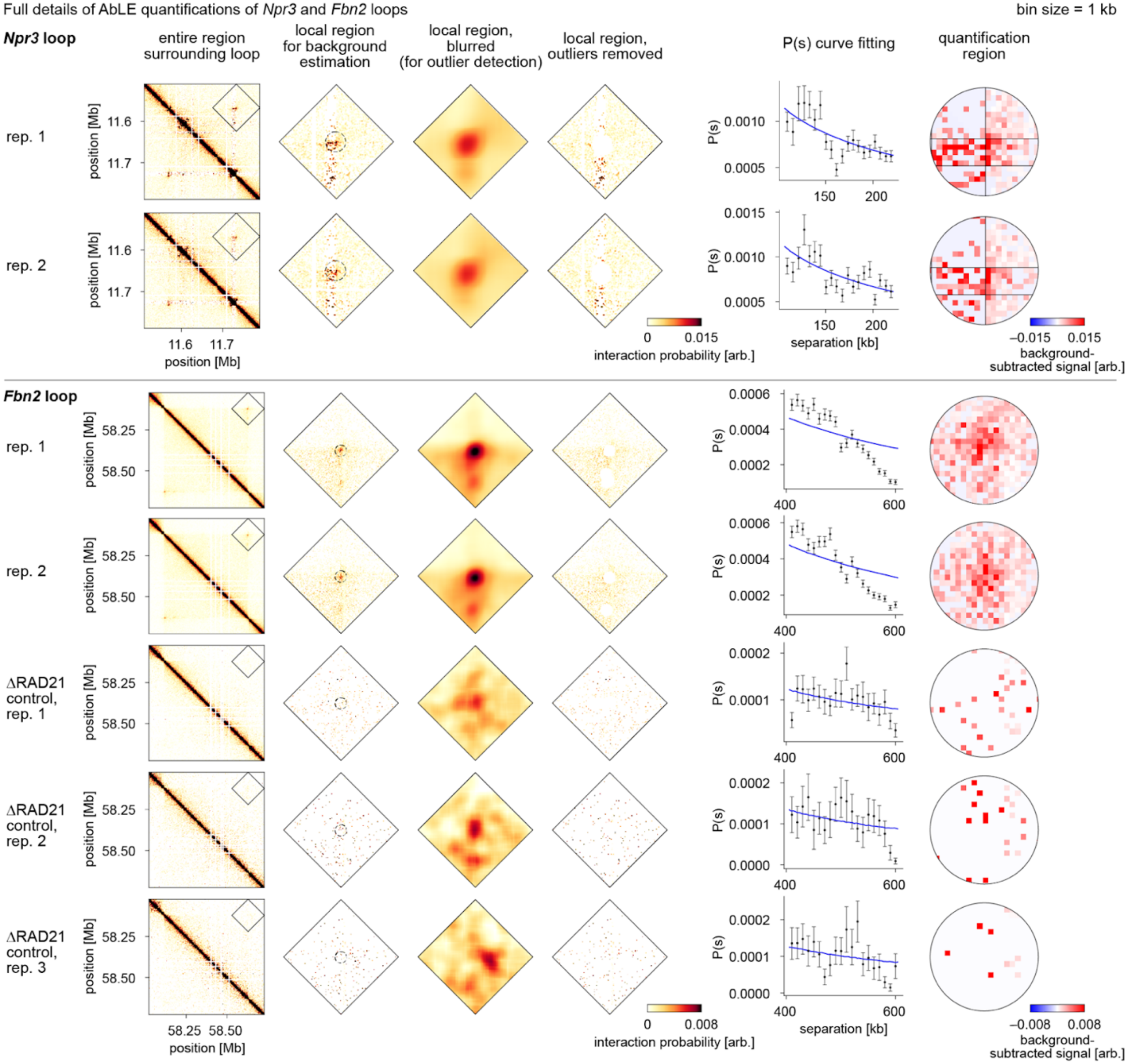


**Figure S4. Full details of AbLE quantifications of *Npr3* and *Fbn2* loops.** For each loop, a diamond-shaped region surrounding the loop is taken. The region is blurred, and all prominent local maxima are identified as outliers. All data points in the region, excluding those within a 10-kb Euclidean distance of outlier points, are used to fit a local $P(s)$ curve, which defines the local background. The AbLE score is defined as the sum of local-background-subtracted pixels within a 10-kb circle surrounding the loop position. All loop quantifications were performed on Micro-C maps at 1-kb resolution. Plotted color scales are linear. Since the *Npr3* loop is heterozygous, additional steps were introduced into the loop quantification procedure. In calculating the AbLE score for the *Npr3* loop, the local-background-subtracted signal inside the synthetic regions was counted once as usual, but the local-background-subtracted signal outside was doubled. The black lines show the bounds of the synthetic regions rounded to the nearest 1 kb; the left vertical line is barely visible because it is nearly tangent to the circular quantification window.

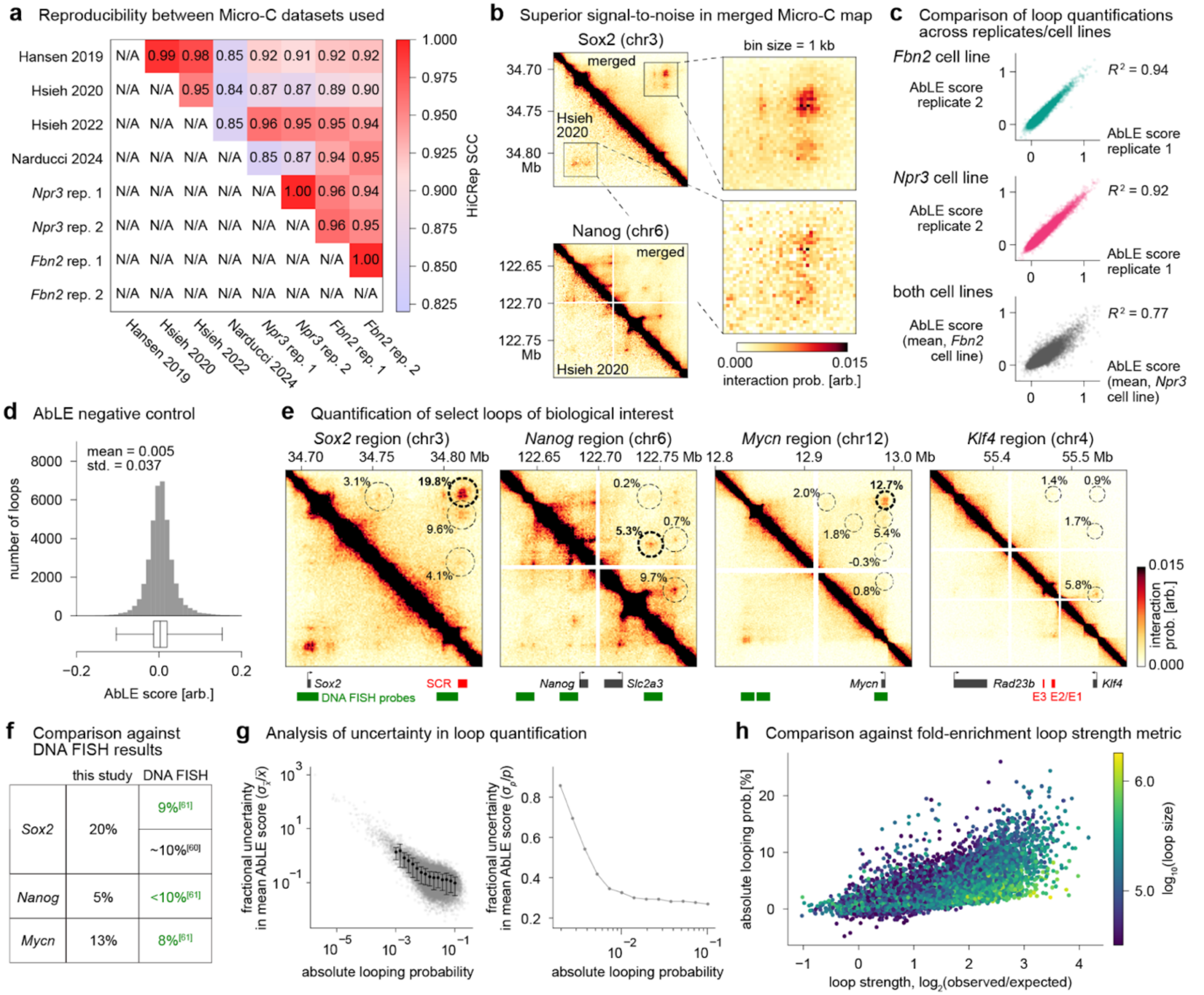


**Figure S5. Supplementary analyses accompanying genome-wide analysis of chromatin looping. a,** Reproducibility between all Micro-C samples used to generate the merged Micro-C map, as quantified by HiCRep[70] stratum-adjusted correlation coefficient (SCC) at 100-kb resolution. **b,** Examples of superior signal-to-noise ratio in the merged Micro-C map compared to the deepest dataset previously available (Hsieh et al., 2020). Micro-C maps are shown at 1-kb resolution with linear color scale; coordinates are in mm39. **c,** Comparison of AbLE scores between replicates in *Fbn2* cell line (top), between replicates in *Npr3* cell line (middle), and between *Fbn2* and *Npr3* cell lines (bottom). **d,** Negative control of AbLE: distribution of AbLE scores for 37,726 "loops" at random locations of the Micro-C map not corresponding to actual loops. Box plot whiskers represent $0.5^{\text{th}}$ and $99.5^{\text{th}}$ percentiles; outliers not shown. **e,** Quantification of select loops of biological interest. In each window, all filtered loops are labeled with their looping probability estimates. Micro-C maps are shown at 1-kb resolution with linear color scale; coordinates are in mm39. Below Micro-C maps, genes are shown in gray, enhancers in red, and DNA FISH probes from Le et al., 2024[61] in green. SCR = *Sox2* control region;[62] E1/E2/E3 = enhancers in *Klf4* region[71]. **f,** Comparison of absolute looping probabilities vs. estimates of contact probability from DNA FISH. DNA FISH contacts were defined as 3D interactions below a threshold of 200 nm.[61] **g,** Uncertainty in absolute looping probabilities arises from biological and technical noise and uncertainty in the calibration factor used to convert to absolute units. Biological and technical noise is captured by the variation in AbLE scores between the four Micro-C replicates (left); the fractional uncertainty (standard error of the mean divided by the mean) is calculated assuming that the samples/replicates are independent measurements. Multiplying the AbLE scores by the calibration factor of 0.186 ± 0.047 (fractional uncertainty = 0.253), the uncertainties are propagated to obtain the uncertainties for the absolute looping probability (right). **h,** Comparison of absolute looping estimates with log-ratio metric for quantifying loop strength.

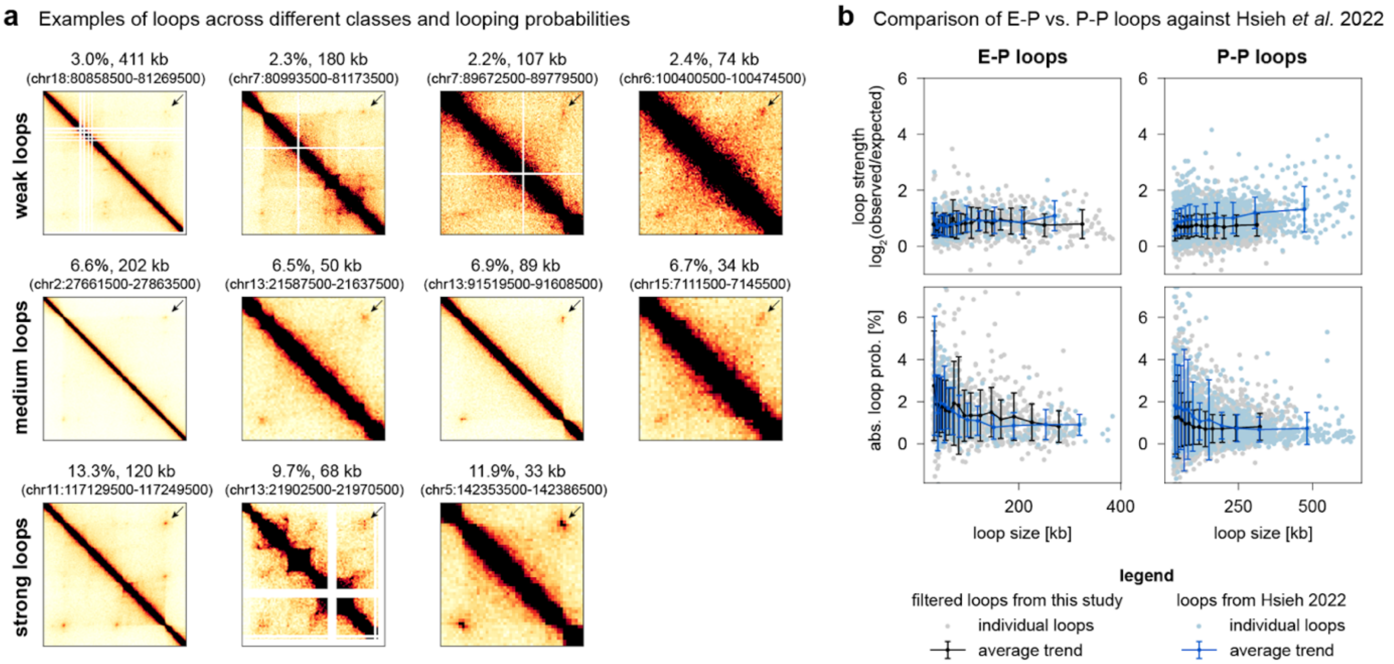


**Figure S6. Supplementary results accompanying looping probability analyses for different loop classes. a,** Examples of weak, medium, and strong loops of different loop classes. Loop classes are defined using the exclusive definitions, i.e., pure CTCF-CTCF loops, pure P-P loops, etc. (no other features allowed). The Micro-C maps shown are an average of the four replicates from the data generated in this study. Micro-C maps are shown at 1-kb resolution with linear color scale. **b,** Comparison of the strength of E-P vs. P-P loops in this study's dataset (black) vs. Hsieh et al. 2022 dataset[46] (blue), using the log-ratio loop strength metric (top) and absolute looping probability (bottom). While the trends in E-P looping were similar, this study found many weak P-P loops compared to Hsieh et al. 2022. These trends are reflected by the log-ratio loop strength metric (used in Hsieh et al. 2022) as well as the absolute looping probability. Error bars indicate the sample standard deviation within each bin.

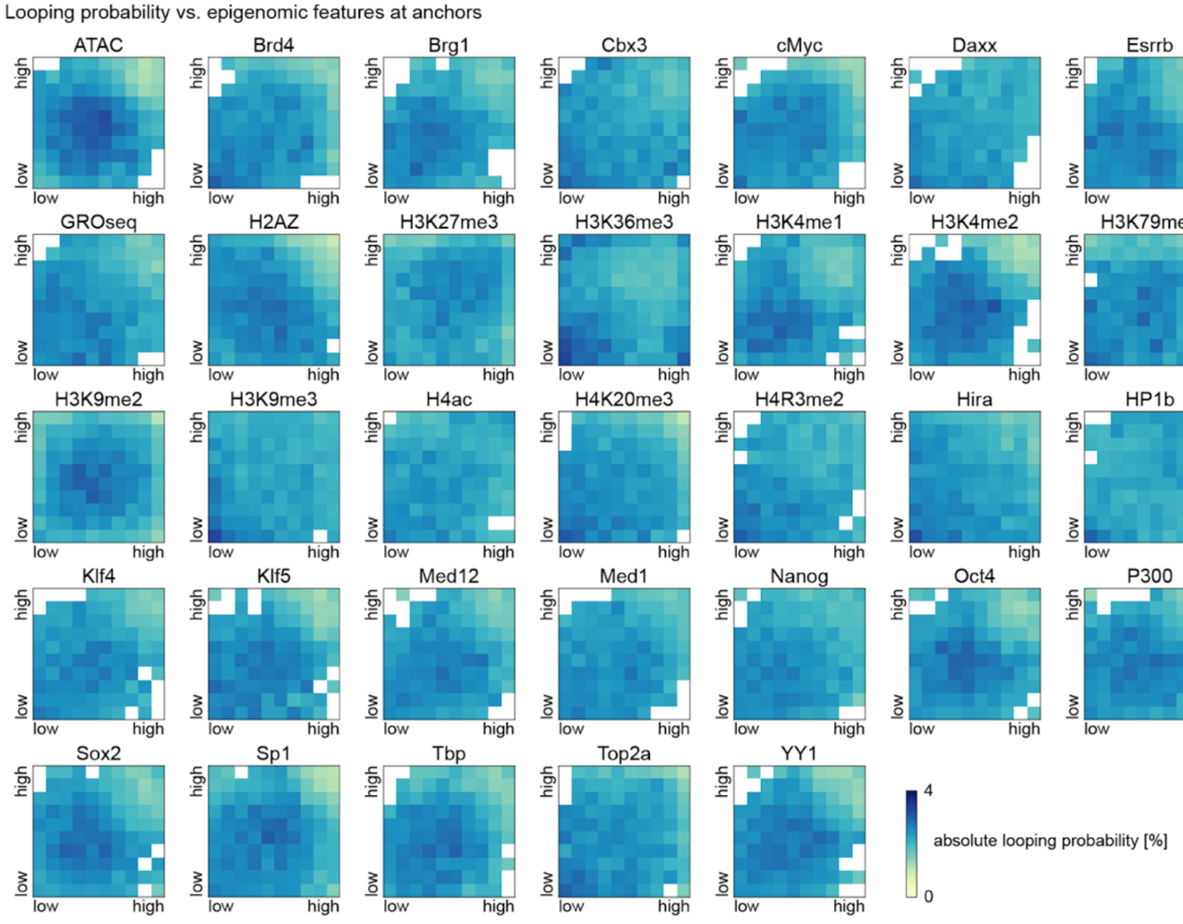


**Figure S7. Looping probability vs. epigenomic feature strengths at loop anchors.** Heatmaps of absolute looping probability vs. feature strengths at loop anchors for all features not shown in **Fig. 3b**. *x*- and *y*- values represent ChIP signals at left and right anchors respectively and are binned into deciles; color scale represents the mean absolute looping probability within each bin. Bins in which the standard error of the mean ($\sigma/n$) exceeds 0.2 are colored white due to their high uncertainty.

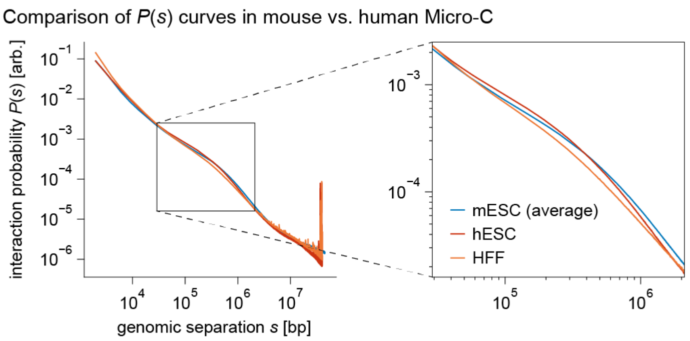


**Figure S8. Comparison of *P*(*s*) curves in mouse vs. human Micro-C.** Plots of genome-wide averaged *P*(*s*) curves in mouse Micro-C (mESC, average across four replicates produced in this study) and human Micro-C (hESC and HFF from Krietenstein et al. 2020[47]). Inset shows *P*(*s*) curve for *s*-values of loops, ranging from 32 kb to 2 Mb.

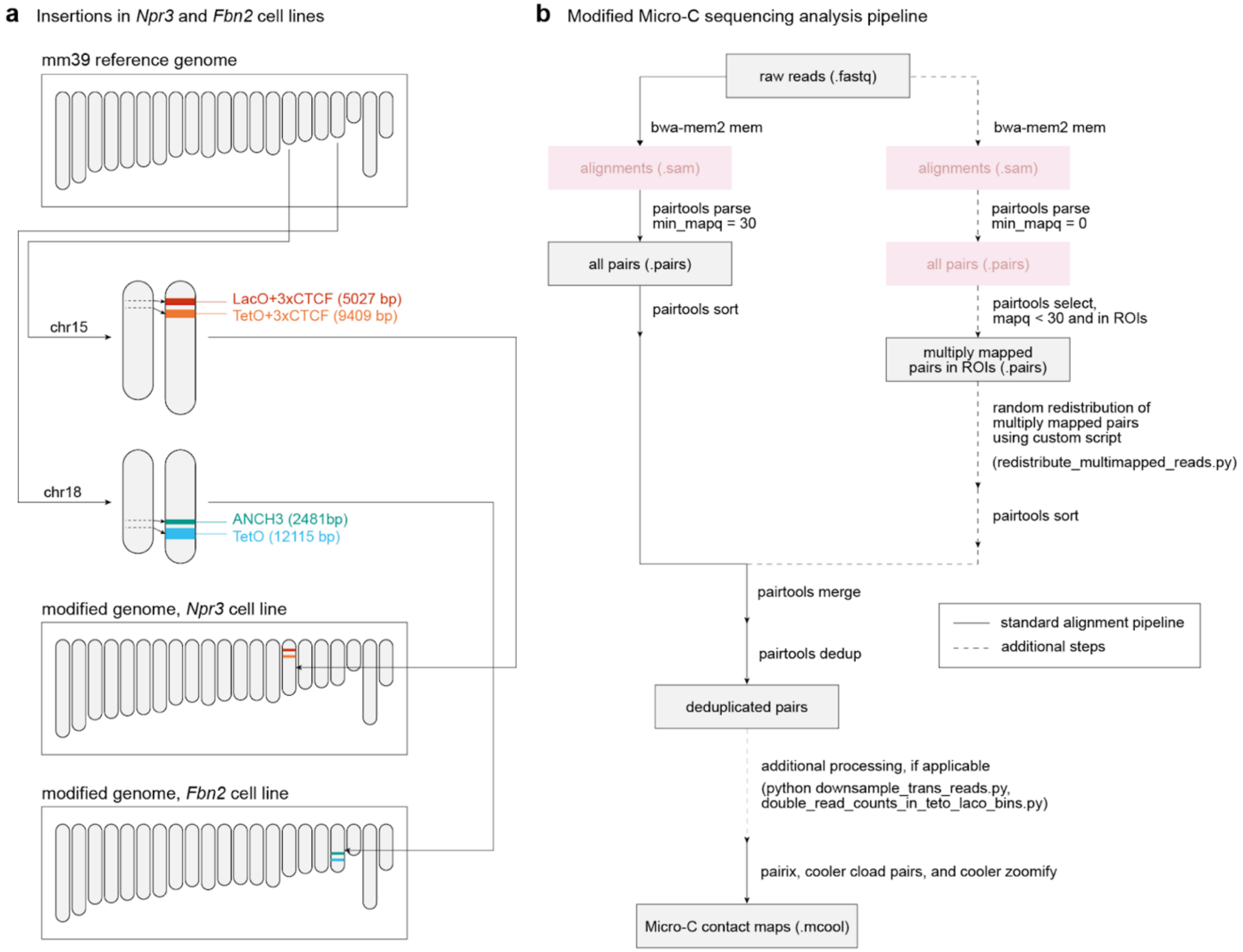


**Figure S9. Additional considerations for processing Micro-C data from cell lines with engineered genomes. a,** Schematic of synthetic insertions in *Npr3* and *Fbn2* cell lines. In the *Npr3* cell line, the insertions included two 3xCTCF binding sites to create the synthetic *Npr3* loop, as well as LacO and TetO arrays to recruit fluorophores near the loop anchors. In the *Fbn2* cell line, ANCH3 and TetO were inserted to recruit fluorophores near the loop anchors of the endogenous *Fbn2* loop. For loop quantification (**Fig. S4**), Micro-C data were aligned to the appropriate modified genomes. **b,** Modified Micro-C sequencing analysis pipeline used in this study. Dotted lines indicate additional custom steps: recovering multiply mapped regions (necessary due to highly repetitive sequences), normalizing the *cis*/*trans* ratio, and doubling the read counts in one of the insertions that was heterozygous.

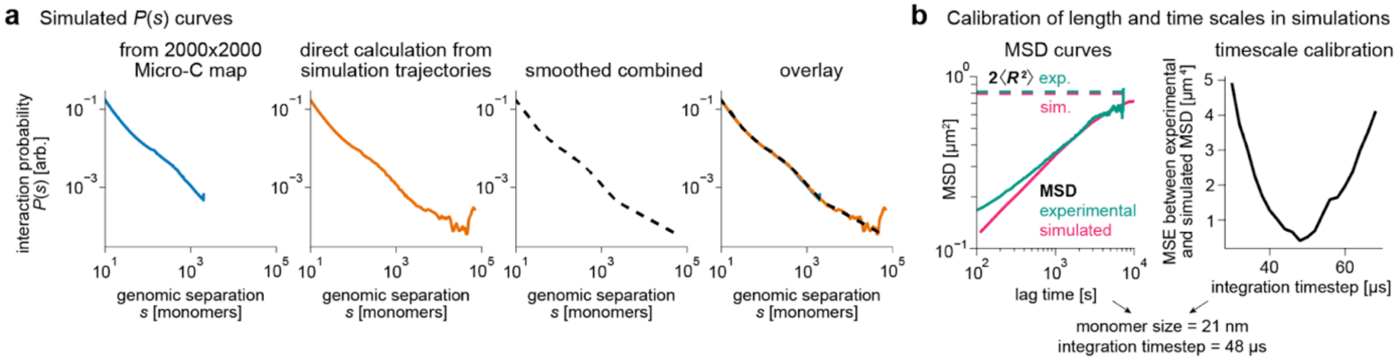


**Figure S10. Additional technical details of 3D polymer simulations. a,** Simulated *P*(*s*) curves from the 3D polymer simulations. The *P*(*s*) curves calculated using two different methods were combined and used to generate a chromosome-wide simulated Micro-C map. **b,** Calibration of length and time scales in simulations to real units. The dotted lines represent the values to which the MSD curves are expected to converge over time. Experimental data are from live imaging of the *Fbn2* loop in the ∆RAD21 condition in Gabriele et al. 2022[20].

### MATERIALS AND METHODS

#### Cell culture

Two mouse embryonic stem cell (mESC) lines were used in this study: clone C36 from Gabriele et al. 2022[20], derived from JM8.N4 cells[72], and the TetO-LacO+3×CTCF cell line (clone 1B1) from Mach et al. 2022[21], derived from E14Tg2a cells[73]. The cell lines had fluorescent labels at the ends of the *Fbn2* and *Npr3* loops, respectively, and are referred to as the *Fbn2* and *Npr3* cell lines here.

*Fbn2* cells were cultured on plates pre-coated with 0.1% gelatin solution (Sigma-Aldrich, G1890) in KnockOut Dulbecco's Modified Eagle Medium (Thermo Fisher, 10829018) supplemented with 15% fetal bovine serum (HyClone, SH30396.03), 2 mM GlutaMAX (Thermo Fisher, 35050061), 1% MEM Non-Essential Amino Acid Solution (Thermo Fisher, 11140050), 0.1 mM 2-mercaptoethanol (Sigma-Aldrich, M3148), 1000 U/mL leukemia inhibitory factor (home-made[74]), 10 μM MEK inhibitor (Tocris, PD0325901), 3 μM GSK3 inhibitor (Sigma-Aldrich, SML1046), and 100 μg/ml Penicillin-Streptomycin (Thermo Fisher, 15140122) in 5% $CO_2$ at 37 °C. Half the medium was replaced every day, and cells were passaged every two days by dissociation with TrypLE Express (Thermo Fisher, 12605028).

*Npr3* cells were cultured on plates pre-coated with 0.1% gelatin solution (Sigma-Aldrich, G2500) in Glasgow Minimum Essential Medium (Sigma-Aldrich, G5154) supplemented with 15% fetal calf serum (Eurobio Abcys), 1% L-Glutamine (Thermo Fisher, 25030024), 1% Sodium Pyruvate MEM (Thermo Fisher, 11360039), 1% MEM Non-Essential Amino Acids (Thermo Fisher, 11140035), 0.1 mM 2-mercaptoethanol (Thermo Fisher, 31350010), 20 U/mL leukemia inhibitory factor (Miltenyi Biotec, premium grade), 1 μM MEK inhibitor (Axon, 1408), and 3 μM GSK3 inhibitor (Axon, 1386) in 8% $CO_2$ at 37 °C. The medium was exchanged daily and cells were passaged every two to three days by dissociation with Accutase (Sigma-Aldrich, A6964).

#### Micro-C experiments and analysis

##### Micro-C experimental procedure

Micro-C was performed on the *Fbn2* and *Npr3* cell lines with two replicates each. The experiment was performed according to a previously published protocol from our laboratory[56] that builds upon the protocol originally developed by Hsieh et al.[45,48–50] Briefly, 5 million cells per replicate were crosslinked for 45 minutes with 3 mM disuccinimidyl glutarate at room temperature. In the final 10 minutes of crosslinking, formaldehyde was added to a final concentration of 1%. After 45 minutes, the reaction was quenched with 0.375 M Tris pH 7.5, and the pellets were snap frozen. At this stage, the *Npr3* cell pellets were shipped on dry ice and remained frozen for 72 hours during transport. Next, the crosslinked chromatin was digested to mononucleosomes using micrococcal nuclease (Worthington Biochemical, LS004798) at a concentration determined by a prior MNase titration. Biotin-dATP and biotin-dCTP were incorporated into the digested chromatin ends in a reaction with 1 U per 1 μg of DNA of Klenow fragment (New England Biolabs, M0210L). The chromatin ends were proximity ligated in a reaction with 20 U/μL T4 DNA ligase (New England Biolabs, M0202M), and the biotin-dNTPs were removed from the unligated ends using 5 U/uL exonuclease III (New England Biolabs, M0206L). Finally, the crosslinking was reversed by incubation overnight at 65 °C in a solution containing 2 mg/mL proteinase K (Viagen Biotech, 501-PK), 10% SDS, 250 mM NaCl, and 0.1 mg/mL RNase A (Thermo Fisher, EN0531). The DNA was subsequently purified using the Zymo DNA Clean and Concentrator (DCC-25) kit (Zymo Research, D4034). The purified DNA was run on a 1% agarose gel, and the di-nucleosomal band was size selected by cutting the band between 250 and 400 bp. The DNA from the band was purified using the Zymoclean Gel DNA Recovery Kit (Zymo Research D4008), and proximity ligated DNA was isolated by streptavidin bead purification with Dynabeads MyOne Streptavidin T1 (Thermo Fisher, 65601). The purified DNA proceeded to library preparation following the NEBNext Ultra II DNA Library Prep Kit for Illumina (NEB, E7645), and the libraries were purified using 0.9X Ampure XP beads (Beckman Coulter, A63881). Library quality was assessed via Fragment Analyzer and qPCR at the MIT BioMicro Center. Pooled libraries were sequenced as 150 bp paired-end reads on an Illumina NovaSeq X system by the Broad Institute of MIT and Harvard's Walk-Up Sequencing services.

##### Initial processing of Micro-C data

Base calls for NovaSeq output were obtained using bcl2fastq (v2.20.0.422), resulting in .fastq files for each sample, pair mate, and flow cell lane. Read quality was verified using FastQC (v0.12.1). Paired-end reads were aligned to mm39-based genomes using bwa-mem2 (v2.2.1) with default settings. Since both cell lines contained insertions of synthetic DNA sequences at the anchors of the loops analyzed in this study, alignment was performed to versions of the mm39 reference genome modified to include these insertions (**Fig. S9a**). Aligned paired-end reads were then parsed with pairtools (v1.0.3)[75] parse with `--add-columns mapq --walks-policy all --min-mapq 30`.

#### Recovery of multiply mapped reads in Micro-C

The synthetic insertions in the *Fbn2* and *Npr3* cell lines contain significant subsequences that are not unique, either because they are internally repetitive (in the case of the TetO arrays) or contain subsequences that also exist elsewhere in the wild-type mouse genome. Conventional approaches for processing Micro-C sequencing data discard all reads that do not map uniquely to a single location, which would result in missing data in the synthetic insertions, preventing accurate quantification of the nearby *Fbn2* and *Npr3* loops. Therefore, additional steps were performed to recover any multiply mapped reads that aligned to the synthetic sequences (**Fig. S9b**). Separately from the main pipeline, aligned paired-end reads were parsed using pairtools parse with `--add-columns mapq,algn_ref_span --walks-policy all --min-mapq 0`. Subsequently, pairtools select was used to filter for pairs where one or both sides had a mapping quality (mapq) below 30 and aligned to a region of interest (defined in **Table S7**). The regions of interest were chosen to include (1) all synthetic sequences that contain internal repetition, (2) all synthetic sequences that appear elsewhere in the wild type genome, and (3) all wild type sequences that appear in the synthetic regions. Only exact matches 35 bp or longer were considered, as Micro-C experiments rarely generate fragments shorter than this length. Under these selection criteria, any pairs containing reads that originated from the synthetic sequence but failed to map uniquely were selected for further processing.

Within the selected pairs, each read that failed to map uniquely was randomly assigned to one of its possible mappings according to the following algorithm. For each side of the pair, all possible hits within the regions of interest were determined. If either side mapped somewhere outside the regions of interest but not uniquely (indicated by mapq < 30), the pair was discarded, as recovering these pairs is beyond the scope of our algorithm. Pairs containing null-mapping reads were also discarded. Next, all pairs formed by all possible hits on one side combined with all possible hits on the other side were considered. If all of these pairs were *trans* (i.e., on different chromosomes), one of the possible pairs was chosen from a uniform distribution. Otherwise, if any of the pairs were *cis* (on the same chromosome), only the *cis* pairs were considered, since *cis* interactions occur with much higher frequency than *trans* interactions. In this case, each *cis* pair was assigned a weight of $s^{-1}$, where $s$ is the genomic distance between the centers of the two reads that form the pair (pairs with $s < 1000$ were assigned a weight of 1/1000), and one of the *cis* pairs was chosen randomly using the normalized weights as the probability distribution. The final result of this custom algorithm is a list of recovered multiply-mapping pairs that have been assigned unique positions.

#### Generating Micro-C maps

For each sample, pairs originating from the standard alignment were merged with the newly recovered multiply-mapping pairs. Duplicate pairs were then removed using pairtools dedup with `--max-mismatch 1`.

The ratio of *cis* to *trans* pairs varies across samples. Since the balancing algorithm for Micro-C matrices takes into account both *cis* and *trans* pairs[76], in order to ensure that quantitative comparisons could be made between different Micro-C maps, we downsampled *trans* pairs to maintain their fraction at a constant 5% of total pairs (**Fig. S9b**). To achieve this, *trans* pairs in the *Fbn2* replicates were retained with probabilities of 0.31 and 0.29, and *trans* pairs in the *Npr3* replicates were retained with probabilities of 0.76 and 0.79.

Next, pairs were indexed using pairix (v0.3.7) and subsequently converted to .cool format using cooler (v0.9.3) cload pairs, creating binned read counts across the genome for 250-bp bins. For the *Npr3* replicates only, all read counts in bins with one or both positions falling in the synthetic regions (with coordinates rounded to the nearest 250 bp) were doubled (**Fig. S9b**). This step ensured that the synthetic *Npr3* loop in the *Npr3* cell line, which is heterozygous, could be compared with the homozygous *Fbn2* loop in the *Fbn2* cell line, as well as all other loops in the genome. Finally, .cool files were converted to .mcool format with cooler zoomify with `--balance --balance-args '--ignore-dist 20000'`, compiling read counts for bins of the following sizes: 250, 500, 1k, 2k, 5k, 10k, 20k, 50k, 100k, 200k, 500k, 1M, 2M, 5M, and 10M bp.

### Calibration of Micro-C dot strength and absolute looping probability

#### Quantification of *Fbn2* and *Npr3* loops with AbLE

We define the AbLE score of a loop as the sum of local-background-subtracted pixel values in the 1-kb resolution Micro-C map, within a circle of radius 10 kb centered on the loop position.

To estimate the local background near a loop, we first obtained a diamond-shaped window on the Micro-C map extending ±$x$ kb from the diagonal, where $x = \sqrt{d/0.0512}$, and $d$ is the size of the loop in kb. This formula for $x$ results in $x = 25$ kb for the smallest loops ($d = 32$ kb), which ensures that the window does not contain the diagonal of the Micro-C map. For loops of other sizes, the window scales with the square root of the loop size, which effectively keeps the fitting error constant (**Fig. S2f**). For imputing missing values and detecting outliers, we used “observed-over-expected” windows, generated by dividing the observed values in a window by their expected values according to the genome-wide average $P(s)$ curve. Any NaN values in the observed-over-expected window were imputed to be the median value in the window. To detect outlier pixels, we applied Gaussian blur with $\sigma = 10$ kb to the observed-over-expected window, and any pixels whose values exceeded two times the median value within the window were considered to be outliers. Finally, to obtain the local background, we fitted the function $b(s) = c \times P(s)$ to all pixels in the original window that were not within 10 kb of an outlier pixel in Euclidean distance. Here, $c$ is an unknown constant that accounts for regional variation in local background, $P(s)$ is the genome-wide averaged $P(s)$ curve for that sample, and $s$ is the genomic separation.

To calculate the AbLE score, a circle of radius 10 kb surrounding the dot was obtained from the Micro-C map at 1-kb resolution, and the previously determined local background was subtracted from all pixels. The AbLE score is defined as the sum of background-subtracted pixel values in the circle. Since the *Npr3* loop is heterozygous, when calculating the AbLE score for this loop only, the local-background-subtracted signal outside the synthetic region was doubled. This is because the Micro-C background-subtracted signal outside the synthetic regions is an average of two alleles, only one of which contains the *Npr3* loop, so after matrix balancing, the signal due to the *Npr3* loop would have effectively been attenuated by a factor of 2. The Micro-C signal inside the synthetic regions could only have been due to the *Npr3* loop and therefore required no modifications.

The details of the AbLE quantifications for the *Fbn2* and *Npr3* loops, including the original windows, blurred windows, $P(s)$ fitting, and circular quantification regions are shown in **Fig. S3.** Additional Micro-C was not performed for the *Fbn2* ΔRAD21 control in this study; instead, existing data of three replicates from Hsieh et al. 2022[46] and Gabriele et al. 2022[20] were used.

#### Bayesian Inference of Looping Dynamics (BILD) on *Fbn2* and *Npr3* loops

To quantify absolute looping probabilities, we applied BILD on past live imaging data of the *Fbn2*[20] and *Npr3* loops[21], using the previously published method and Python code [20]. Briefly, BILD defines two states ("looped" and "unlooped") from control experiments and then segments the WT/untreated data into those states using a hierarchical Bayesian model. While the "unlooped" control is simply the ΔCTCF condition, we obtain a proxy for the "looped" state by rescaling the mean squared distance $\langle R^2 \rangle$ observed in the ΔRAD21 condition with the fraction of genomic distance remaining between the two loci when the loop is formed (17 kb for *Fbn2*, 6.5 kb for *Npr3*). From these calibration data, BILD then assembles a Rouse polymer model to represent these two states: in the "unlooped" state the model is a linear chain of $3L+1$ monomers, with the two loci of interest positioned at monomers $L+1$ and $2L+1$; we parametrize this model by maximum posterior fitting to the ΔCTCF data. For the "looped" state we insert an additional bond between the two loci of interest, such that the effective spatial distance between them decreases to the calibration value for the looped state. This two-state Rouse model then provides the likelihood function needed for the hierarchical Bayesian segmentation, which is performed as described in the original publication.

#### Determining constant of proportionality between AbLE score and absolute looping probability

To determine the proportionality constant between AbLE score and absolute looping probability, we fitted a line through the origin ($y = kx$ where $k$ is the unknown parameter) to two data points, corresponding to the *Fbn2* and *Npr3* loops. We performed an orthogonal distance regression to fit the line using `scipy.odr`[77]. The $x$-value of each point was taken as the mean of the AbLE scores for the two replicates, and the error in the $x$-direction was calculated as the sample standard deviation of the AbLE scores: $\sqrt{(x_1 - \bar{x})^2 + (x_2 - \bar{x})^2}$, where $\bar{x} = (x_1 + x_2)/2$. The $y$-value of each point was taken as the mean of 1,000 bootstrap estimates of the absolute looping probability from BILD, and the error in the $y$-direction was taken as the standard deviation of these estimates. The *Fbn2* ΔRAD21 control point was not used for fitting.

### Analyses of chromatin looping probability in mESCs

#### Merging all existing mESC Micro-C datasets

A merged Micro-C map was generated by combining the Micro-C data from this study with existing mESC Micro-C data from previous studies (**Table S2**). Raw sequencing reads from all four samples generated in this study were aligned to the wild-type mm39 reference genome using bwa-mem2 (v2.2.1) with default settings. Raw reads from previous studies were downloaded from the National Center for Biotechnology Information's (NCBI) Sequence Read Archive (SRA) using the SRA Toolkit (v3.0.10) and aligned to the mm39 reference genome with default settings. Aligned reads were parsed and deduplicated in each sample separately. Pairs from all samples were merged into a single file using pairtools merge, and this file was indexed, converted to .cool format, and finally converted to .mcool format. All steps used the same software and settings as described in the preceding sections.

#### Calculation of chromosome-averaged *P*(*s*) curves

For each sample, a *P*(*s*) curve was calculated from the Micro-C map to obtain the relationship between interaction probability and genomic separation, which is necessary for downstream analysis (loop filtering and quantification). The curve was calculated for each chromosome individually using the `expected_cis` function from cooltools (v0.5.4) on a Micro-C map with 1-kb bin size. Then, *P*(*s*) curves were averaged across chromosomes 1-19 and chrX, weighted by the chromosome size in bp, resulting in one representative genome-wide *P*(*s*) curve for each sample.

#### Chromatin loop calling with Mustache

Chromatin loops were identified genome-wide from the merged Micro-C map with 1-, 2-, and 5- kb bin sizes using *Mustache* (v1.0.1).[59] The sparsity threshold was set to 0.7 for 1-kb bins and 0.88 at 2- and 5-kb bins, and the q-value threshold was set to 0.1. The genomic intervals corresponding to the loop anchors were expanded bidirectionally by 5-kb using bedtools (v2.27.1) slop, and any two loops identified at two different bin sizes but with overlapping anchors were declared to be duplicates, as identified by bedtools pairToPair. For duplicate loops, the position defined with the smallest available bin size was taken as the final position.

#### Filtering of quantifiable chromatin loops

Chromatin loops were filtered based on their suitability for quantification according to several characteristics. Any loops whose anchors were less than 32 kb apart were discarded, as the diagonal of the Micro-C map would interfere with the measurement of the loop. This resulted in 140,830 loops among the original 153,658 loops. Next, loops with either anchor within 5 kb of a stripe of NaN values in the balanced map were discarded, resulting in 121,656 loops. For each loop, the Micro-C map with 1-kb bin size was obtained on a square window extending ±10 kb from the loop position. Only loops with greater than 0.4 reads per pixel (176 reads) in this window in all four samples were retained, resulting in 60,716 loops. Next, we sought to filter out cases where multiple loops were within the same window, since this complicates accurate quantification. For these loops, for the visualization window in the merged Micro-C map was considered: each pixel value in the window was divided by its expected value as calculated from a genome-wide *P(s)* curve, any NaN values were imputed as the mean value within the window, and a Gaussian blur with $\sigma = 2.5$ px was applied. Loops whose global maxima existed within a Euclidean distance of 2.5 px of the center pixel were retained, since a global maximum elsewhere implies that any interference from a stronger loop nearby or that the loop is extremely weak. These filtering steps resulted in a final list of 36,804 quantifiable loops across the mESC genome.

#### Quantification of chromatin loops genome-wide

For each of the filtered loops, the absolute looping probability was estimated by calculating the AbLE score in all four Micro-C samples, averaging across samples, and multiplying by 0.186, the scaling factor determined from calibration with live imaging data.

AbLE scores for filtered loops were calculated using the same procedure as for the *Fbn2* and *Npr3* loops, but the merged Micro-C dataset was used for the outlier detection step.

#### Calculation of relative loop strength using log-ratio metric

The following procedure was used to calculate relative strength of loops using the log-ratio metric, i.e., $\log_2$(observed/expected): a 41x41 viewing window centered on the loop was obtained from the Micro-C map of bin size = 1 kb. The observed signal was calculated as the average value in the 5×5 square at the center of the window. The expected signal was calculated as the average value in the 5×5 squares at the top left and bottom right corners of the window, since these have equivalent genomic separations as the loop. The relative loop strength was defined as the base-2 logarithm of the quotient of the observed signal divided by the expected signal.

#### AbLE negative control at random positions

As a negative control, the AbLE score was calculated for random pairs of loci that did not correspond to actual loops. First, 200,000 random pairs throughout the genome were chosen with the same distribution of genomic separations as the original set of 36,904 filtered and quantified loops. These pairs were filtered using the same criteria as the original loops, except the global maximum step was omitted because no global maximum was expected. The AbLE scores were then calculated for these resulting 37,726 loops.

#### Classification of chromatin loops by mechanism

To classify chromatin loops by mechanism, we first identified whether each anchor overlapped a promoter, enhancer, or CTCF- and cohesin-bound site. Promoter regions were defined using all TSS locations in the mm39 UCSC RefGene annotation[78] ±2 kb. Enhancers were defined based on overlap of H3K4me1 (GSE90893) and H3K27ac (GSE90893) that did not overlap promoters. CTCF- and cohesin-bound sites were defined based on overlap of CTCF (GSE90994) and SMC1A (GSE123636) and the presence of the CTCF binding motif.

For all datasets, raw reads were downloaded from NCBI SRA using the SRA Toolkit and aligned to the mm39 reference genome using bowtie2 (v2.2.5) with default settings, and only reads which aligned uniquely were retained for downstream analysis. Duplicates were removed using sambamba (v1.0.0) markdup. For experiments with multiple replicates, the bam files of the replicates were merged using sambamba merge. Peaks were called using MACS2 (v2.2.9.1) callpeak with `-q 0.01`. The `--broad` flag was used for H3K4me1 and H3K27ac to call broad peaks, whereas narrow peaks were called by default for CTCF and SMC1A.

CTCF peaks were then overlapped with CTCF motifs identified using FIMO (v5.4.1): first, fasta-get-markov was used to generate a background model using the mm39 genome assembly, then motifs were identified using --max-stored-scores 50000000 --thresh 1e-3. Only the motif with the highest score within each CTCF peak was retained. Lastly, these motif positions were intersected with SMC1A peaks using bedtools intersect (v2.30.0) to obtain a final list of CTCF- and cohesin-bound sites. Meanwhile, H3K4me1 and H3K27ac peaks were intersected using bedtools intersect to obtain enhancer sites.

Loop anchors were classified as promoters, enhancers, or CTCF and cohesin-bound if they overlapped any of these features within ±2.5 kb. Since many anchors contained more than one feature (e.g., both an enhancer and CTCF/cohesin binding), we classified loops using both an "inclusive" scheme and an "exclusive" scheme. Under the inclusive definitions, P-P loops included all loops with promoters at both anchors, E-P loops included all loops with an enhancer at one anchor and a promoter at the other anchor, E-E loops included all loops with enhancers at both anchors, and CTCF-CTCF loops included all loops with CTCF- and cohesin-bound sites at both anchors such that the CTCF motifs were in a convergent orientation. Any anchor that did not overlap a promoter, enhancer, or CTCF- and cohesin-bound site was classified as "other". Under the exclusive definitions, only loops with exactly one feature at each anchor were considered; e.g., P-P refers to all loops with promoters at both anchors, but no enhancers or CTCF- and cohesin-bound sites at either anchor.

#### Quantification of epigenomic features at loop anchors

All public epigenomic datasets (**Table S4**), with the exception of GRO-seq (GSE69143), were downloaded as raw reads from NCBI SRA using the SRA Toolkit and aligned to the mm39 reference genome using bowtie2 with default settings. Adapters were trimmed automatically upon download, except for Daxx (GSE70811), for which adapters failed to be trimmed automatically and were instead trimmed manually using cutadapt (v4.6). Duplicates were removed using sambamba markdup. For experiments with multiple replicates, the bam files of the replicates were merged using sambamba merge. Next, bedtools genomecov was used to generate pileups in bedgraph format, and the scale parameter was set such that the resulting output coverage was given in reads per million. The bedgraph files were converted to bigwig files using UCSC bedGraphToBigWig.[63] The signal of each epigenomic feature, defined as the sum of values of the bigwig at all bases over a 5-kb interval centered on the loop anchor, was calculated at the left and right anchors of all filtered and quantified chromatin loops.

For GRO-seq (GSE69143), bigwig files for the positive and negative strands were downloaded from the NCBI Gene Expression Omnibus (GEO), merged, and converted to bedgraph files using bigWigToBedGraph and lifted over from mm9 to mm39 coordinates using liftOver. Overlapping intervals were removed, and all values were scaled down by a factor of 0.02, which was chosen arbitrarily but results in a scale that is comparable to other datasets. The resulting bedGraph was converted to bigWig, and signal at loop anchors was calculated as described above.

#### Linear models for predicting looping probabilities from epigenomic features at loop anchors

In our linear model of absolute looping probability as a function of epigenomic feature strengths, each of the 36,804 filtered and quantified loops was taken as a single data point. The signals of each feature at the left and right anchors, plus loop size, were used as regressors ($x$-values, 87 variables total), and the absolute looping probability was the dependent variable ($y$-value). Loops with any $x$- or $y$-values less than or equal to zero were removed. All $x$- and $y$-values were then log-transformed and subsequently scaled to have zero mean and unit variance, and loops with any $x$- or $y$-values greater than 4 or less than –4 (i.e., further than 4 standard deviations from the mean) were removed. This resulted in 29,822 loops used for downstream analysis. Linear models were trained on a randomly chosen subset consisting of 80% of loops (23,858 loops). Due to symmetry between the left and right anchors, we doubled the number of data points in the training set by creating 23,858 new points by swapping the left and right anchor values for each feature.

To implement the ordinary least squares (OLS) model, we used `linear_model.LinearRegression` from scikit-learn.[79] We applied projection to latent structures (PLS) using `cross_decomposition.PLSRegression` from scikit-learn to reduce the dimensionality of the input space. We took the first four components from PLS, which was sufficient to achieve nearly the same $R^2$ as the OLS model. Similar to principal components analysis (PCA), each component in PLS is a linear combination of the original features. To increase the interpretability of the results, we performed a varimax rotation of the first four components from PLS, resulting in four new components spanning the same space. We considered all subsets of these four rotated components, finding that just two of these components alone was sufficient to achieve nearly the same $R^2$ as the OLS model. These components were used in our final linear model and subsequently denoted by $z_1$ and $z_2$.

#### Analysis of chromatin looping probabilities from human Micro-C data

The hESC and HFF Micro-C datasets were analyzed using the same pipeline and parameters used on the mESC Micro-C data. Briefly, raw data was downloaded from NCBI SRA using the SRA Toolkit, aligned to the hg38 reference genome using bwa-mem2, and pairs were processed using the same steps as the Micro-C data in this study, including downsampling *trans* pairs to achieve a constant *trans* fraction of 0.05. *Mustache* was used to call loops on each dataset, and loops were filtered for quantifiability based on various criteria. For each loop, the AbLE score was calculated and multiplied by 0.186 to obtain an estimate of the absolute looping probability. Loops whose left and right anchor positions were within 10 kb in hESCs and HFFs were identified as shared loops.

We identified a small number of loops with very high looping probabilities in both hESCs and HFFs. Upon visual inspection, these extreme outliers appeared to be generated by anomalous patterns in the Micro-C contact maps that were likely technical artifacts. We manually identified 16 regions containing these patterns (**Table S8**), and we excluded all loops whose left and right anchors were within these regions from all analyses (82 loops in hESCs, 102 loops in HFFs, and 35 shared loops). These loops constituted a very small fraction of the total loops.

CTCF- and cohesin-bound sites in hESCs were identified by intersecting called peaks from CTCF ChIP-seq (ENCSR000AMF) and RAD21 ChIP-seq (ENCSR000ECE). CTCF- and cohesin-bound sites in HFFs were identified by intersecting called peaks from CTCF ChIP-seq (ENCSR000ECE) and SMC1A CUT&Tag (4DNEXMHZPZVI). Loop anchors were defined as CTCF sites if they overlapped CTCF sites within ±2.5 kb. All analysis was performed in hg38 coordinates. Shared hESC/HFF CTCF-CTCF loops were defined as those found in both cell types and identified as CTCF-CTCF loops in both cell types.

#### 3D polymer simulations

##### Design of simulated chromosome

We performed 3D polymer simulations of a single chain consisting of $G$ = 70,000 monomers, with each monomer corresponding to 1 kb of DNA. This chain represented a fictitious chromosome consisting of 35 identical repeats of a 2 Mb region (2000 monomers), with individual monomers representing CTCF sites or enhancer/promoter sites. Within each repeat of 2000 monomers, the locations and orientations of these sites were as follows:

- Left-pointing CTCF positions: [574, 694, 866, 1241, 1390, 1580, 1752, 1800]
- Right-pointing CTCF positions: [200, 330, 724, 1425, 1433, 1604]
- Enhancer/promoter positions: [250, 372, 540, 745, 775, 833, 961, 1202, 1330, 1640, 1722]

The region was designed by hand to generate looping interactions that span a wide range of sizes, reflecting typical loop sizes observed in real chromatin.

##### 1D simulations of loop extrusion

To model the effect of loop extrusion by cohesin, we applied a 1D simulation as described in previous work.[80] A cohesin contains two motor subunits, and these subunits were programmed to move bidirectionally along the polymer, moving away from each other one monomer at a time. Also, cohesins were not allowed to bypass each other or move past the first and the last monomers on the chromosome. Cohesins could be stalled upon encountering CTCF sites in a convergent direction, with a stalling probability intrinsic to the CTCF site. The relative cohesin stalling probabilities at the CTCF sites were as follows:

- Relative cohesin stalling probabilities by left-pointing CTCFs: [0.6, 0.8 , 0.95, 0.1 , 0.6, 0.6, 0.8, 0.1]
- Relative cohesin stalling probabilities by right-pointing CTCFs: [0.9, 0.3 , 0.95, 0.4 , 0.3, 0.4]

Once one of cohesin's two motor subunits became stalled at a CTCF site, that subunit was no longer allowed to move until the entire cohesin dissociated from the DNA. Meanwhile, the other motor subunit of the cohesin was allowed to continue extruding until it was either stalled at another CTCF site or until the entire cohesin dissociated from the DNA.

A fixed number of cohesin molecules was used for the 1D loop extrusion simulations. The number of cohesins was determined by the ratio of chromosome length to the average separation *d* between cohesins (i.e., the inverse of the density). When a cohesin dissociated from the DNA, it was immediately reloaded at another pair of adjacent monomers with uniform probability over the entire chromosome, excluding monomers that were occupied by existing cohesin motor subunits. The dissociation rate of cohesin was a function of the processivity $\lambda$ (i.e., the average length of DNA extruded by an unobstructed cohesin before it dissociates) and the stabilization of cohesin stalled at CTCF sites, modeled as a fold-increase *b* in cohesin lifetime. In this simulation we used $d = 240$ monomers, $\lambda = 300$ monomers, and $b = 4$.

#### 3D polymer simulations via OpenMM

We performed a 3D simulation to model the molecular dynamics of the chromosome. The simulation was performed using the *Polychrom* package (v0.0.1),[81] which wraps the OpenMM molecular simulation toolkit.[82] In our model, cohesin positions were informed by the results of a parallel-running 1D simulation, and cohesins act as harmonic bonds connecting the two monomers occupied by their motor subunits. The 1D cohesin extrusion simulations were run for a total of 10,000 translocation steps in order to reach a steady state before coupling them to the 3D simulation.

In the 3D simulation, all adjacent pairs of monomers experienced a harmonic potential (`polychrom.forces.harmonic_bonds`) that equals zero at the equilibrium length of 1 and increases to kBT at a displacement of 0.1. Adjacent pairs of monomers were also subject to a harmonic angle force (`polychrom.forces.angle_force`) whose energy at 1 radian was equal to 0.05kBT. Furthermore, all monomers in the chain exhibited an interaction potential with all other monomers defined by the smooth square well potential (`polychrom.forces.heteropolymer_SSW`), accounting for volume exclusion and monomer attraction effects simultaneously. The repulsion energy was set to 3kBT, allowing for some chain crossings, and the repulsion radius was set to 1.05 monomer units. To model enhancer/promoter interaction, there was also an attractive potential of 3kBT between any of the monomers designated as enhancer/promoter sites.

At the start of each simulation, the polymer was initialized in a compact loop on a cubic lattice (`polychrom.starting_conformations.grow_cubic`) with a volume number density of 30% in terms of the monomer length scale. The system thermostat was set with an error tolerance of 0.01, and the collision rate was set to 1. Throughout the simulation, the polymer was subject to a spherical confinement force (`polychrom.forces.spherical_confinement`) with $k = 1$ and whose size was chosen to maintain the 30% density. The simulation was run for a total of 1,000,000 blocks, each consisting of 2,360 Langevin dynamics polymer integration steps, and blocks ranging from 500,000 to 1,000,000 were used for the analysis, sampled every 100 blocks.

#### Calculation of dot strength in simulated Micro-C maps

A simulated Micro-C map of the 2000-monomer repeating region was generated using Polychrom's `monomerResolutionContactMapSubchains` function with a cutoff of $r_{\mu C} = 3$ units (in **Fig. 1d**,**e**) or $r_{\mu C} = 2$, 3, or 5 units (in **Fig. S2**) , i.e., any pair of monomers less than $r_{\mu C}$ units apart was counted as a contact. The $P(s)$ curve for $s \leq 2000$ monomers was then calculated using the data in this contact map. Meanwhile, another $P(s)$ curve was calculated for $10 \leq s \leq 70{,}000$ (the length of the entire chromosome) directly from the simulation trajectories: for a given value of *s*, the probability of being less than $r_{\mu C}$ units apart was calculated among all pairs of monomers with a separation *s* along the polymer. As expected, these two $P(s)$ curves were approximately the same for values of *s* less than 2,000. To create a smoothed version of the $P(s)$ curve, we fitted a 6th-degree polynomial to $P(s)$ vs. $\log(s)$ for $s \leq 2000$ and a linear function to $P(s)$ vs. $\log(s)$ for $2000 \leq s \leq 30{,}000$ and manually set the values of $P(s)$ for $2000 \leq s \leq 2908$ to connect the two portions. The $P(s)$ curves described here are shown in **Fig. S10a**. We then generated a simulated Micro-C map of the entire 70,000-monomer chromosome, where the contact probability within each region (2000×2000 blocks along the diagonal) was taken directly from the original regional contact matrix, and the contact probability across different regions was taken to be the value of the smoothed $P(s)$ curve at the separation *s* of the bin coordinates. The simulated Micro-C map was exported in .cool format using cooler load and balanced using cooler balance `--ignore-dist 20000`, which were the same settings used to process the experimental Micro-C maps.

#### Calculation of ground-truth looping probabilities in simulations

The ground-truth looping probability of a given CTCF-CTCF loop in the simulation was defined as the fraction of the 35 repeats of the loop that was held together by cohesin(s) at a given time, averaged over all time points used for the analysis: blocks 500,000 to 1,000,000, sampled every 1000 blocks. It was possible for a loop to be held together by more than one cohesin. For example, if a cohesin molecule was located between positions *X* and *Y*, and another cohesin molecule was located between *X* and *Z*, then positions *Y* and *Z* would also be considered as looped.

The ground-truth looping probability of enhancer/promoter loops in the simulation was calculated as the fraction of the 35 repeats of the loop such that both ends were within $r_{EP}$ = 3 units (**Fig. 1e**) or $r_{EP}$ = 2, 3, or 5 units (**Fig. S2**) in 3D space, averaged over blocks 500,000 to 1,000,000 sampled every 1000 blocks, minus the background looping probability. The background looping probability for any two monomers was calculated from another simulation that was run without enhancer/promoter attraction but contained all other features including loop extrusion.

#### Calibration of simulation length and time scales

The length scale and time scale of the 3D simulation were calibrated by analyzing the curve of mean squared distance (MSD) vs. time. The MSD is formally defined as the mean value of the squared magnitude of the change in the distance vector between a pair of monomers over a lag time $\Delta t$; the average is taken over all pairs of time points separated by $\Delta t$. We performed a simulation of our chain with no loop extrusion and no enhancer/promoter attraction, while keeping other parameters to be the same as in the original simulation. The resulting MSD curve was calculated between two monomers with a separation of 515 monomers, such that it could be fitted against the actual experimental MSD curve for the *Fbn2* loop as measured using live imaging in Gabriele et al. 2022[20] (**Fig. S10b**). We first matched the $2\langle R^2 \rangle$ values in the experimental and simulated data to obtain the monomer size, ≈21 nm. Then, we minimized the mean squared error between the experimental and simulated MSD curves to estimate the integration timestep to be ≈48 µs (duration of each block ≈0.11 s, entire simulation ≈30 h). Taking into account the 1D simulation parameters described above, the resultant cohesin extrusion speed was 550 bp/s per motor subunit.

### SUPPLEMENTARY TABLES

**Tables S1-S6.**

- **Table S1.** List of mESC Micro-C datasets used to create merged Micro-C map
- **Table S2.** List of original loop calls from Mustache
- **Table S3.** List of filtered loop calls from Mustache with loop quantifications and classifications
- **Table S4.** List of public epigenomics datasets used in analysis
- **Table S5.** Quantifications of epigenomic feature signals at left and right anchors of all filtered mESC loops
- **Table S6.** List of filtered loop calls from Mustache in hESC and HFF Micro-C data from Krietenstein et al. 2020, with loop quantifications and classifications

Tables S1-S6 are available at the following Dropbox link:
https://www.dropbox.com/scl/fo/ynl853n8jn5jcxidxwr4s/AOFAvVMcJMwUg_AvOxX92YY?rlkey=rrah3j7c4sql6j8pd0hhp5ys1&e=1&st=v5svyxc5&dl=0

**Table S7.** Regions of interest for recovery of multiply mapped reads. Coordinates are reported in the modified mm39 genomes that contain the synthetic insertions, using the 1-based indexing convention.

| name | bounds of region | description |
|---|---|---|
| *Npr3* cell line | | |
| A | chr15:11567091-11572418 | 3xCTCF+LacO insertion |
| B | chr15:11722583-11732291 | TetO+3xCTCF insertion |
| C | chr1:34257393-34257803 | sequences elsewhere in the genome that are identical to a section of the TetO+3xCTCF and LacO+3xCTCF insertions |
| D | chr4:132977929-132978340 | |
| E | chr8:13511840-13512242 | |
| *Fbn2* cell line | | |
| F | chr18:58104053-58116467 | TetO+Hygro insertion |
| G | chrX:105230186-105230991 | sequences elsewhere in the genome that are identical to a section of the TetO+Hygro insertion |
| H | chr12:10950107-10950463 | |

**Table S8.** Regions of hg38 reference genome excluded from analysis of hESC/HFF Micro-C data from Krietenstein et al. 2020[47] due to potential artifacts in Micro-C map.

| chrom | start | end | chrom | start | end |
|---|---|---|---|---|---|
| chr1 | 12850000 | 13400000 | chr10 | 4900000 | 5080000 |
| chr1 | 25250000 | 25400000 | chr12 | 131050000 | 131750000 |
| chr4 | 143750000 | 144150000 | chr14 | 105500000 | 105800000 |
| chr5 | 650000 | 900000 | chr15 | 44800000 | 45100000 |
| chr6 | 31220000 | 31550000 | chr16 | 2540000 | 2700000 |
| chr6 | 32450000 | 32800000 | chr16 | 20400000 | 20630000 |
| chr6 | 167150000 | 167420000 | chr22 | 25200000 | 25700000 |
| chr7 | 48830000 | 48930000 | chr22 | 42480000 | 42600000 |
